# Genomic diversity and global distribution of four new prasinoviruses from the tropical North Pacific

**DOI:** 10.1101/2024.10.02.616384

**Authors:** Anamica Bedi de Silva, Shawn W. Polson, Christopher R. Schvarcz, Grieg F. Steward, Kyle F. Edwards

## Abstract

Viruses that infect phytoplankton are an integral part of marine ecosystems, but the vast majority of viral diversity remains uncultivated. Here we introduce four near-complete genomic assemblies of viruses that infect the widespread marine picoeukaryote *Micromonas commoda*, doubling the number of published genomes of *Micromonas* dsDNA viruses. All host and virus isolates were obtained from tropical waters of the North Pacific, a first for viruses infecting green algae in the order Mamiellales. Genome length of the new isolates ranges from 205-212 kb, and phylogenetic analysis shows that all four are members of the genus *Prasinovirus*. Three of the viruses form a clade that is adjacent to previously sequenced *Micromonas* viruses, while the fourth virus is relatively divergent from previously sequenced prasinoviruses. We identified 61 putative genes not previously found in prasinoviruses, including a phosphate transporter from a family novel to viruses, and a potential apoptosis inhibitor novel to marine viruses. Forty-eight genes in the new viruses are also found in host genome(s) and may have been acquired through horizontal gene transfer. By analyzing the coding sequences of all published prasinoviruses we found that ∼25% of prasinovirus gene content is significantly correlated with host genus identity (i.e., *Micromonas*, *Ostreococcus*, or *Bathycoccus*), and the functions of these genes suggest that much of the viral life cycle is differentially adapted to the three host genera. Mapping of metagenomic reads from global survey data indicates that one of the new isolates, McV-SA1, is relatively common in multiple ocean basins.

**Importance statement:** The genomes analyzed here represent the first viruses from the tropical North Pacific that infect the abundant phytoplankton order Mamiellales. Comparing isolates from the same location demonstrates high genomic diversity among viruses that co-occur and presumably compete for hosts. Comparing all published prasinovirus genomes highlights gene functions that are likely associated with adaptation to different host genera. Metagenomic data indicates these viruses are globally distributed, and one of the novel isolates may be among the most abundant marine viruses.

## Introduction

A significant fraction of marine viral diversity is composed of viruses that infect phytoplankton, the diverse unicellular primary producers that perform the majority of marine photosynthesis (Suttle, 2007). Viruses that infect phytoplankton affect biogeochemical processes by lysing their hosts, thereby shunting nutrients and energy to smaller microbial cells, and by shaping phytoplankton production through mortality, metabolic manipulation, and virus-driven trait evolution (Wilhelm and Suttle, 1999; Weitz and Wilhelm, 2012; Hurwitz *et al*., 2013; Våge *et al*., 2013).

In this study we focus on prasinoviruses, which are double-stranded DNA viruses that infect prasinophytes (Weynberg 2017; ICTV 2023). Prasinophytes are a diverse and paraphyletic group of green algae (Leliaert et al., 2012), and the *Prasinovirus* genus is made up of double-stranded DNA viruses in the family *Phycodnaviridae*. However, the isolated prasinoviruses all infect members of the phytoplankton order Mamiellales, which are included under the generic term prasinophyte (Bachy et al., 2021). This order of algae includes the three cosmopolitan genera *Bathycoccus*, *Ostreococcus*, and *Micromonas* (Yau et al., 2015; ICTV, 2023). The Mamiellales are ubiquitous in the sunlit ocean and are often major community members in both oligotrophic and eutrophic environments, typically dominating the picoeukaryotic fraction of primary producers under productive conditions (Not et al., 2004; Lopes Dos Santos et al., 2017). These algae are known for their small size, which is exemplified by the species *Ostreococcus tauri*, considered to be the smallest free-living eukaryote at ∼0.8 µm cell diameter, while *Bathycoccus* and *Micromonas* are ∼2 µm in diameter (Manton and Parke, 1960; Eikrem and Throndsen, 1990; Courties et al., 1994). The first isolated virus infecting eukaryotic microalgae was a *Micromonas* virus (Mayer and Taylor 1979), and isolates of prasinoviruses have been used to study many topics, such as viral production and decay (Cottrell and Suttle, 1995), marine viral gene content (e.g., Finke et al., 2017; Bachy et al., 2021), viral alteration of host nutrient uptake (Monier et al., 2017), consequences of host resistance to lytic infection (e.g., Thomas et al., 2012; Heath et al., 2017), and diel changes to the dynamics of viral infection (e.g., Derelle et al., 2015). Prasinophytes and their viruses are useful model systems for understanding the biology and functional consequences of phytoplankton viruses because they are cosmopolitan, relatively abundant, and amenability to laboratory manipulation (Moreau et al., 2010; Clerissi et al., 2014; Lopes Dos Santos et al., 2017). In environmental samples from the surface ocean, prasinoviruses are typically among the most abundant members of the *Nucleocytoviricota*, which is a highly diverse phylum of large, eukaryote-infecting dsDNA viruses (Farzad et al., 2022; Ha et al., 2023). As of this writing, 22 genomes of prasinovirus isolates have been published, including four *Micromonas* viruses, five *Bathycoccus* viruses, and 12 *Ostreococcus* viruses, and prior studies of have described key characteristics of prasinovirus gene content (Moreau et al., 2010; Finke et al., 2017; Bachy et al., 2021). Genes shared among all published prasinovirus genomes (i.e., core genes) are largely associated with basic viral functions, such as DNA replication, transcription, and nucleotide metabolism (Moreau et al., 2010). In contrast, non-core genes (i.e., genes present in a clade but not shared by all members) include many involved in cellular metabolism, such as nutrient acquisition, photosynthesis, and carbohydrate metabolism (Moreau et al., 2010; Finke et al., 2017). These genes associated with diverse cellular processes are presumed to alter host metabolism to enhance viral replication, but although few prasinovirus genes have been experimentally evaluated for function (Monier et al., 2017).

The characterization of additional viral isolates and their hosts continues to be important, as isolates contribute to the database of viral reference genomes with known host taxa, while also allowing experimental studies of the ecology and (co)evolution of host-virus systems. Sequencing additional prasinovirus representatives would facilitate a better understanding of the scope of prasinovirus genetic diversity, as well as how genome content has diverged among prasinovirus clades. In the current work we sequence and analyze the genomes of four *Micromonas commoda* viruses isolated from coastal and open-ocean waters around the island of Oahu, Hawaiʻi. These four isolates represent the first prasinovirus genome assemblies from tropical Pacific waters, and they double the number of published *Micromonas* phycodnavirus genomes. Our analysis of these genomes has four aims: (1) establish the phylogenetic relationships of the new isolates to other prasinoviruses; (2) evaluate the gene content of the new isolates in relation to other prasinoviruses and two *Micromonas* hosts to characterize gene novelty, potential gene origins, and genomic diversity among co-occurring prasinoviruses with overlapping host range; (3) quantitatively compare gene content across all prasinovirus genomes to understand what processes might underlie adaptation of viruses to different host genera; and (4) evaluate the distribution of the new isolates in the global ocean using metagenomic surveys.

## Methods

### Virus isolation

Four virus strains infecting the marine eukaryote *Micromonas commoda* were examined in this study. Three of the strains, McV-KB2, McV-KB3, and McV-KB4, were isolated from the surface waters (< 2 m) of Kāne‘ohe Bay (21°27′N 157°48′W) on the windward side of O‘ahu (Schvarcz 2018). The fourth strain, McV-SA1, which was previously referred to as MsV-SA1 in Schvarcz (2018), was isolated from a depth of 25 m from the pelagic research site Station ALOHA (22°45’N 158°W). For simplicity, we will refer to this suite of virus strains as “HiMcVs”, an abbreviation of Hawai‘i *Micromonas commoda* Viruses. The four HiMcVs overlap in host range, based on lysis tests with seven *Micromonas* strains isolated from Kāne‘ohe Bay (UHM1060–UHM1065) and Station ALOHA (UHM1080), with each viral strain infecting 2–6 *Micromonas* strains (Fig. 1). All virus strains are maintained in the UHM culture collection via propagation on their original hosts, which were isolated from the same waters as their corresponding virus strains. Full isolation methods are described in Schvarcz (2018). In brief, whole sea water from respective sites was filtered, concentrated, and then added to healthy cultures which were subsequently monitored for lysis. If lytic effects were confirmed after multiple transfers to healthy culture, lysates were further purified through several rounds of dilution-to-extinction. In the current study, lysate stocks were maintained through fortnightly transfers of lysate into healthy cultures grown in f/2-Si medium (Guillard and Ryther, 1962; Guillard, 1975). Once lysis took place, typically within 4-6 days of the initial challenge, lysates were stored at 4°C.

**Figure 1.**
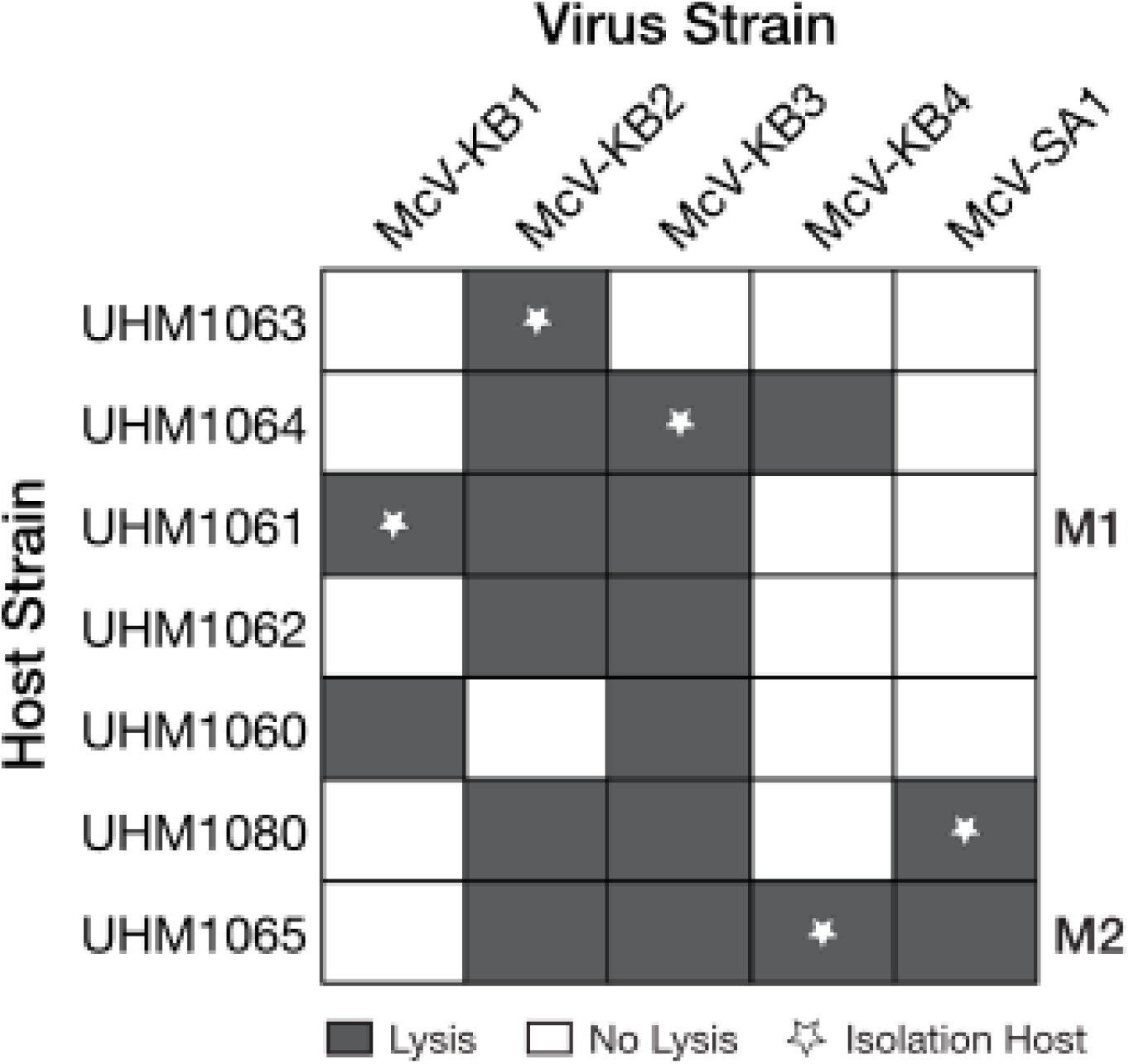
Host range of virus strains used in this study. UHM culture collection IDs are listed for 7 *Micromonas* isolates across the top, with Micromonas host strains listed on the left-hand side. Shortened names for cell strains UHM1061 and UHM1065 are listed on the right-hand side. Dark gray squares indicate l successful lytic infection. Stars represent the host from which each virus strain was isolated.

### Electron microscopy

Virions in the McV-SA1 lysate were purified by banding in a cesium chloride equilibrium buoyant density gradient. The virion-containing fraction was buffer exchanged into SM using a centrifugal ultrafilter and a drop adsorbed onto a carbon-stabilized formvar on a 200-mesh copper grid. The grid was rendered hydrophilic by glow discharge within a few hours prior to sample deposition. Virions were negatively stained with uranyl acetate and then visualized by transmission electron microscopy (Hitachi HT7700).

### Whole-genome sequencing

Purification, extraction, and sequencing protocols for McV-SA1 are described in Schvarcz (2018). In summary, McV-SA1 virions were purified by banding in a CsCl density gradient as noted above for electron microscopy. DNA was extracted from purified virions with a MasterPure™ Complete DNA and RNA purification Kit (Epicentre Biotechnologies, now Biosearch Technologies). PacBio library preparation and sequencing (P6-C4 chemistry) for a pooled sample of DNA from distantly related algal viruses, including McV-SA1, was performed with Sequel II SMRT cells at the University of Washington PacBio Sequencing Services facility. McV-SA1 reads were extracted using high-similarity BLAST searches against references assemblies. Illumina NextSeq 150bp paired-end sequencing was conducted in tandem at the Georgia Genomics and Bioinformatics Core at the University of Georgia, USA. The isolates McV-KB2, McV-KB3, and McV-KB4 were prepared without using a density gradient and sequenced by Illumina short-read technology. Prior to generating lysates for these viruses, we used a combination of filtration and antibiotics to reduce contamination from bacteria and phage present in the host cultures. *Micromonas* host cultures were filtered onto a 1 µm track-etched polycarbonate membrane filter (Nuclepore, Whatman) and then resuspended into sterile f/2 -Si medium containing a cocktail of broad-spectrum antibiotics (Supplementary Table S1). Cultures were transferred three to four times into fresh antibiotic-containing f/2 -Si at 1:100 inoculum:medium. Flow cytometry counts indicated a tenfold reduction of bacteria in treated *Micromonas* cultures. To obtain McV DNA for sequencing, 5 µL of 0.2 µm-filtered lysate was added to 50 mL of the antibiotic-treated host culture. After lysis, cultures were filtered through a 0.2 µm pore cellulose acetate syringe filter, and subsequently concentrated by centrifugal ultrafiltration in units with 10 kDa MWCO regenerated cellulose membranes (Amicon^®^ Ultra, Millipore Sigma). DNA was then extracted from concentrated samples (Wizard^®^ Genomic DNA Purification kit, Promega). Library preparation and sequencing for Illumina 151bp paired-end reads were performed at SeqCenter (formerly Microbial Genome Sequencing Center), located at the University of Pittsburgh, USA.

### Genome assembly and annotation

Genome assembly methods varied among the viruses depending on sequencing method. PacBio reads for Station ALOHA strain McV-SA1, extracted from a multiplexed sample through BLAST, were assembled with Canu v.1.0 (Koren *et al*., 2017) and polished with NextSeq data using Quiver v2.0.0, with 100% agreement with Illumina reads after polishing (Schvarcz, 2018). The Kāne‘ohe Bay virus strains (McV-KB2, McV-KB3, and McV-KB4) were assembled using two approaches that created relatively complete assemblies. The first approach, used for McV-KB3, used the default assembler in Geneious 11.1.5. Illumina 151bp paired-end data were trimmed of adapters (kmer = 27), and low-quality reads (minimum = Q20) and short reads (minimum = 20bp) were excluded using the BBDuk plug-in ((v1.0; Biomatters Ltd.) and then normalized with the Geneious built-in tool (default settings) before assembly. We performed a nucleotide sequence similarity search of the resulting contigs with the Basic Local Alignment Search Tool (BLASTn; Altschul *et al*. 1990) against the NCBI non-redundant (nr) nucleotide database (Sayers *et al*. 2022) and identified two contigs (147 and 60 kb) with similarity to known prasinoviruses. The reads mapped to these contigs were dissolved and re-assembled. This produced a single 205 kb contig. For McV-KB2, the iterative mapping approach using the Geneious assembler did not produce a single contig, so we instead used the metagenome assembly tool metaviralSPAdes (Antipov *et al*., 2020, Galaxy Version 3.15.4+galaxy 2 using SPAdes 3.15.3 accessed through usegalaxy.org) which produced a single 210 kb contig. Finally, for McV-KB4 we were unable to obtain a single contig using either method, but five putative prasinovirus contigs totaling 212 kb were obtained from the Geneious assembler, which we treat as a draft assembly for use in comparative analyses. Illumina raw reads were mapped back to McV-KB2, McV-KB3, and McV-KB4 to check for assembly errors, resulting in 100x to 1000x continuous coverage. We treat the McV-SA1, McV-KB2, and McV-KB3 assemblies as near-complete based on their lengths, which are comparable to known prasinoviruses (Table 1), and based on whole-genome alignments using progressiveMauve, which indicated high synteny of McV-SA1, McV-KB3, McV-KB4, and the most similar previously sequenced *Micromonas virus* isolate, MpV1 (GenBank NC_014767.1; See Results).

**Table 1.**
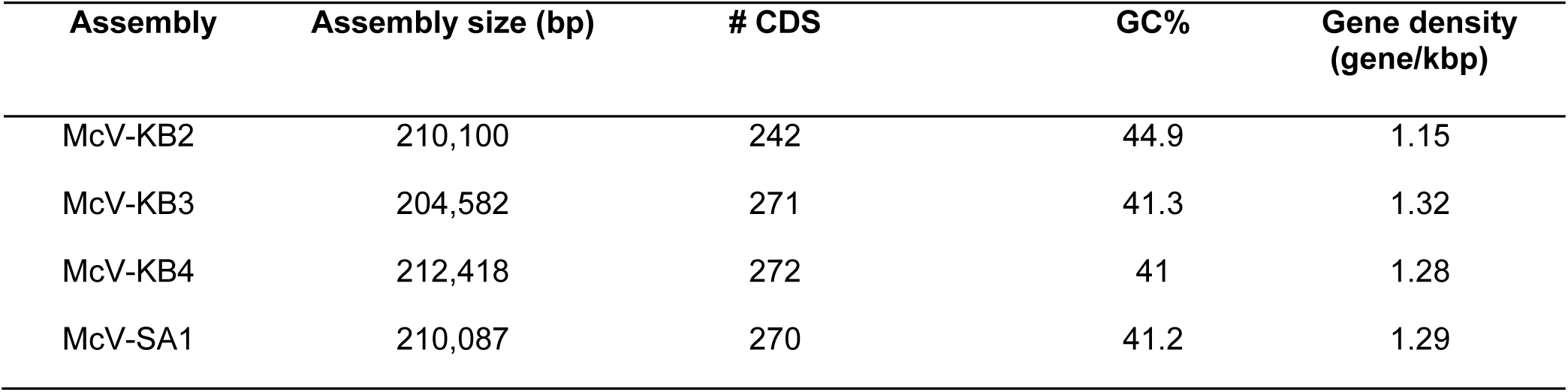
Characteristics of genome assemblies of four Hawaiʻi *Micromonas commoda* virus strains.

| Assembly | Assembly size (bp) | # CDS | GC% | Gene density<br>(gene/kbp) |
| --- | --- | --- | --- | --- |
| McV-KB2 | 210,100 | 242 | 44.9 | 1.15 |
| McV-KB3 | 204,582 | 271 | 41.3 | 1.32 |
| McV-KB4 | 212,418 | 272 | 41 | 1.28 |
| McV-SA1 | 210,087 | 270 | 41.2 | 1.29 |

Gene prediction was conducted with Prokka v.1.14.5 (Seemann, 2014) accessed via Kbase (kbase.us) for all four assemblies. Only CDS with a start and stop codon and a minimum of 65 amino acids (195 nucleotides) were used in downstream analyses.

Functional and structural annotation was performed manually by integrating information from EggNOG mapper (v.1.2, evalue ≤ 0.01, minimum 25% nucleotide identity), the InterProScan web interface (v.5.66-98.0, default settings), and refseq_protein (BLASTp, evalue ≤ 0.001) (Altschul et al., 1990; Quevillon et al., 2005; Cantalapiedra et al., 2021). Additionally, we identified top BLASTp hits against a custom database of genes from the Global Ocean Eukaryotic Virus (GOEV) (Gaïa et al., 2023). We also searched for giant virus MAGs closely related to our isolates by using the isolate genomes as nucleotide BLAST queries against a custom database comprising the GOEV GVMAG contigs. GOEV includes 581 metagenome assembled genomes from uncultured marine viruses and 224 *Nucleocytoviricota* isolate reference genomes.

### Host strain information

The genomes of two *Micromonas* strains in our collection, UHM1061 and UHM1065 (referred to here as M1 and M2 for simplicity), were sequenced. Both strains were isolated from the surface water of Kāne‘ohe Bay. M1 and M2 are lysed by both McV-KB2 and McV-KB3, while M2 is also lysed by McV-KB4 and McV-SA1 (Fig. 1). The 18S rRNA gene sequences of the M1 and M2 cell lines group into a clade with sequences of *Micromonas commoda* (Supplementary Figure S1) and we therefore identify these strains as *Micromonas commoda* (Bedi de Silva *et al*. 2024).

We created high-fidelity reference genomes from M1 (JBHGVY000000000) and M2 (JBHKAF000000000) using PacBio consensus long read (CLR) technology. As our algal cultures were not axenic, we used serial antibiotic treatments (antibiotic recipe in Supplementary Table S1) followed by banding in a continuous Percoll density gradient to obtain a partially purified *Micromonas* fraction. Cells were then pelleted, flash frozen in liquid nitrogen, and shipped to the University of Delaware Sequencing and Genotyping Center. High-molecular-weight DNA was extracted using a modified CTAB method. Library preparation and sequencing was performed with PacBio Sequel II SMRT cells.

A library of transcript sequences was also generated to aid in gene calling. Total RNA was generated from M1 and M2 cell cultures by sampling 10 mL of exponential phase culture (∼10^6^ cell mL^-1^) every four hours for 24 hours to capture diel variation in gene expression. An additional culture sample was given a heat shock treatment to stimulate stress response, in which 10 mL aliquots were placed in a 30°C water bath for 30 minutes. Samples for RNA extraction were syringe filtered onto 25 mm diameter, 1-µm pore size polycarbonate filters (Sterlitech) and then frozen immediately in liquid nitrogen and stored at −80°C. Within a week of sampling, filters were thawed over ice and total RNA extracted using the ZymoBIOMICS RNA Miniprep Kit (Zymo Research). Extracted RNA was sent to University of Delaware Sequencing and Genotyping Center for 81bp paired-end sequencing on the Illumina NextSeq 550.

PacBio CLR data was assembled using Canu (ver 1.9) in PacBio-raw CLR assembly mode, providing an estimated genome size parameter of 22 Mbp (Koren et al., 2017). Resulting contigs were further polished to remove remaining InDel errors by iterative rounds of mapping CLR reads to reference contigs using BLASR (ver 5.3.3 with default parameters except: maxMatch=30, minSubreadLength=750, minAlnLength=750, minPctSimilarity=70, minPctAccuracy=70, hitPolicy=randombest), followed by error correction using Arrow (Pacific Biosciences GenomicConsensus ver 2.3.3 with default parameters except: minCoverage=5, minConfidence=40, coverage=120) until a stable reference was obtained (4 iterations; Chaisson and Tesler, 2012). Closing of circular/organellar elements was performed using Circlator (v1.5.5) and additional manual finishing was performed including manual assessment and scaffolding/overlap of adjacent contigs and resolution/dereplication of haplotype bubbles. Chromosome assignments were manually made using alignment to reference genomes with ProgressiveMauve (v2.4.0) (Worden et al., 2009; GenBank CP001323).

Annotation was performed with Maker (v3.01.03). A custom repeat library was generated using RepeatModeler (v2.0.1) with RepeatScout (v1.0.6) and TRF (v4.0.9). These repeats and repeats for order Mamiellales (CONS-Dfam_3.1-rb20170127) were identified by RMBLAST (v2.10.0) in RepeatMasker (v4.1.0) and masked for gene model annotation. Genome-specific *ab initio* gene calls by GeneMarkES (v2.5p) and SNAP (v2006.07.28) were used to train Augustus (v3.3.3) gene models using e-training scripts (BUSCO v4.0.2).

Illumina RNAseq data was quality trimmed with TrimGalore! (v0.6.5) using Cutadapt (v3.3) and mapped to draft genomes with STAR (v2.7.9a) using the two-pass method and was the basis for a genome-guided transcriptome assembly using Trinity (v2.13.2). Trinity transcripts and primary CDS and protein sequences annotated in *Micromonas pusilla* assemblies RCC299_229_v3.0 and CCMP1545_v3.0, and additional protein sequences extracted from Uniprot for order Mamiellales were mapped to the genome as evidence and used to assess support for *ab initio* gene models.

Noncoding RNA was identified using tRNAscan-SE (v2.0.5) and RNAmmer (v1.2), functional annotations were made by using BLASTp (v 2.12.0) against a Swiss-Prot database (v2021.04), and additional annotations were applied using InterProScan (v5.53-87.0).

### Gene content comparisons

We compared the gene content of the four HiMcVs, to determine 1) how gene content varies among four viruses that overlap in host range and were either isolated from the same coastal location (Kāne‘ohe Bay) or an offshore site (Station ALOHA), 2) which genes are shared between the HiMcVs and the two Hawai‘i *Micromonas commoda* cell strains with sequenced genomes, 3) whether there are genes in our virus strains not seen in previously published prasinoviruses genomes, and 4) whether there are genes that consistently distinguish viruses infecting the three Mamiellales genera (*Micromonas*, *Ostreococcus*, and *Bathycoccus*) that may provide insight into the process of adaptation to different host taxa.

To make these comparisons, we used OrthoFinder (v5.5.; Emms and Kelly, 2019) to identify orthologous groups of genes in a dataset containing the four HiMcVs, the two *Micromonas commoda* strains isolated from O‘ahu, and all 21 previously known prasinovirus genomes published in GenBank. Additionally, we followed Bachy et al. (2021) by including chloroviruses, which form a monophyletic group with prasinoviruses based on marker gene analysis (Bellec et al., 2009) and serve as an outgroup in phylogenetic analyses. Our full genome dataset (accession numbers in Supplementary Table S2) therefore includes strains that infect the Mamiellales genera of *Micromonas* (n = 8, including the four HiMcVs), *Ostreococcus* (n = 12), and *Bathycoccus* (n = 5); *Paramecium bursaria Chlorella* virus strains (n = 4); and the *Micromonas commoda* host strains M1 and M2 (n = 2).

To determine whether viruses infecting the three Mamiellales genera have consistently different gene content, we used gene count data for all prasinovirus genomes and fit the following linear model for each orthogroup:

lm(*orthogroup gene count per genome ∼ host genus*).

This model quantifies the correlation between the number of genes in an orthogroup and the genus of the host infected by a viral strain, to identify orthogroups that most strongly differentiate viruses infecting different host genera. Linear models were compared to a null model using Chi-squared likelihood ratio tests in R (v.4.30; Core Team, 2022). P-values were adjusted for the false discovery rate using the R p.adjust function. Prasinovirus orthogroup count data was also used to create a clustered heatmap using the R pheatmap package (Kolde, 2019; Fig. 6).

### Species and gene tree construction

Phylogenetic relationships of the HiMcV isolates to other members of the order *Algavirales* were assessed by constructing species trees using orthologous genes from the aforementioned published genomes of prasinoviruses and chloroviruses. We used a gene alignment concatenation approach that included all orthogroups shared among all prasinovirus and chlorovirus genomes (i.e. “core” orthogroups; n = 26). We separately aligned each orthogroup using MAFFT (v7.450; Katoh and Standley, 2013), accessed through Geneious. Gene alignments were trimmed to eliminate non-overlapping sequences with Goalign v0.3.7 (https://github.com/evolbioinfo/goalign). If more than one gene copy was present in a genome, the paralog most closely related to orthologs in other genomes was chosen, and then the 26 ortholog alignments were concatenated in Geneious. A phylogeny was estimated with FastTree (v.1.2; Price et al., 2010) within Geneious. We also constructed a tree using only polB sequences, in order to compare the gene tree of this common prasinovirus marker gene to our core gene-based species trees. We used FigTree (v.1.4.4; http://tree.bio.ed.ac.uk/software/figtree/) to visualize both gene concatenation and polB trees (Fig. 3 & Supplementary Figure S2).

OrthoFinder automatically generates a species tree using shared orthogroups with the STAG algorithm (Emms and Kelly, 2018), but this tree does not include node support values when < 100 core orthogroups are present, as is the case with our dataset. We have provided this tree in the supplement as a point of comparison (Supplementary Figure S3).

We identified a core HiMcV alternative oxidase gene that may have recent cellular or phage origins based on the taxonomy of top BLAST hits. We constructed a gene tree to see if the HiMcV orthologs of this gene were more closely related to host or phage sequences, and whether alternative oxidases have been acquired more than once by phytoplankton viruses. We created a MAFFT alignment using alternative oxidase sequences from the four HiMcVs, the two Hawai’i *Micromonas* hosts, OtV RT-2011 (the only other prasinovirus with an alternative oxidase gene), and representative cyanobacteria and cyanophages. This alignment was then trimmed with Goalign and a phylogeny was constructed in FastTree. Previous work on plant alternative oxidases has found that the mitochondrial alternative oxidase (AOX, also called ubiquinol oxidase) and the plastid alternative oxidase (PTOX, also called plastoquinol terminal oxidase) are often misannotated or annotated in an inconsistent way (Nobre et al., 2016). Therefore, we also included reference PTOX and AOX plant and algal sequences to aid in interpreting the alignment and phylogeny.

### HiMcV detection in metagenomes

To assess the global distribution of the four HiMcVs we searched publicly available metagenomes, spanning Pacific and Atlantic ocean basins, from the Hawai‘i Ocean Time-series (HOT), GEOTRACES, and Tara Oceans (Mende et al., 2017; Biller et al., 2018; Brum et al. 2015; Pesant et al., 2015). The HOT data contains samples from 19 timepoints and nine discrete depths above 200 m at Station ALOHA. The other databases contain one sample per station, taken at a depth above 150 m. Sequencing runs from HOT include 20 L samples filtered onto a 0.2 µm filter (n = 293), and samples filtered onto a 0.02 µm filter after passing through at 0.2 µm filter (n = 185). The GEOTRACES metagenomic dataset consists of 100mL samples of whole seawater from the photic zone filtered onto 0.2 µm filters (n = 490), with samples taken from the GA02 and GA03 transects in the North Atlantic, the GA10 transect in the South Atlantic (off the coast of South Africa), and the GP13 transect in the South Pacific (off the coasts of Australia and New Zealand). Metagenome samples from the Tara Oceans Virome database were collected from 100 L whole seawater obtained from 5–100 m depth, depending on the station. Seawater was prefiltered through either a 20 µm or 5 µm net, then put through a 0.22 µm pore size membrane filter. Iron flocculation was then used to concentrate viruses in the filtrate. The metagenomic samples used in our analysis thus include some that should be enriched for free virions in the McV size range (i.e., the fraction between 0.02 and 0.2 µm), and others that should contain primarily cells and larger virions (i.e., the > 0.2 µm fraction), while potentially including smaller virions that did not pass through the 0.2 µm filter. Samples of this larger size fraction were included because a previous study of the GEOTRACES metagenomes (> 0.2 µm) found that putative prasinoviruses were among the most abundant and diverse *Nucleocytoviricota* representatives in these samples (Ha et al., 2023).

We used CoverM v0.6.1 (https://github.com/wwood/CoverM) to search the metagenomic datasets for sequences that mapped onto at least one of the four HiMcV assemblies. Requirements of 95% minimum read identity and 20% minimum covered fraction were used (Ha et al. 2023), indicated with the flags --min-read-percent-identity and --min-covered-fraction. For each successful hit, relative abundances of each virus (i.e., percent of reads from the metagenome sample) derived from CoverM results were then merged with metadata to create a map of hits in R statistical software. CoverM search information, including accession numbers and metadata, are available in Supplementary Table S4.

During the annotation phase of our genomic analysis, we found a metagenome assembled genome from the GOEV database that was similar to McV-SA1. This MAG was derived from TARA ocean data and was the eighth most common assembly in the GOEV database. To provide some context for the presence McV-SA1 in the world ocean, we mapped the reads of the GOEV MAG against the McV-SA1 assembly.

## Results

### Genome assemblies and virus characteristics

Four HiMcV draft genomes were assembled, and we posit these assemblies are near-complete based on nucleotide length, the number of predicted genes, and whole-genome alignments comparing these genomes to each other and to the most similar previously sequenced *Micromonas* virus. The assemblies range from 205 to 212 kbp, with the largest genome belonging to McV-KB4 (Table 1). The McV-KB2 (GenBank Accession: PP911589) genome G+C content (44.9%) was notably higher than the other three genomes (41–41.3%) and its coding density lower (1.15 vs.1.28–1.32 gene/kbp). To help assess genome completeness we created a Mauve alignment with the published genome of MpV1 (NC_014767.1), the virus most closely related to McV-KB3 (PQ109088), McV-KB4 (PQ359806) and McV-SA1 (as described in the next section, Phylogeny). These four genomes exhibited a high degree of synteny, although MpV1 is shorter in total by 20 to 26 kbp (Fig. 2A). It appears that MpV1 has one major ∼11 kbp inversion, relative to the other three genomes, towards the center of its genome. A Mauve alignment using only McV-KB3, McV-KB4 and McV-SA1 showed that these three genomes have high structural similarity (Fig. 2B). Inclusion of McV-KB2 in Mauve alignments resulted in a large number of Locally Collinear Blocks (LCBs), represented as colored blocks in Fig. 2C, which indicated that there were substantial differences in genome organization between McV-KB2 and the other HiMcVs. Overall, the organization of the HiMcV genomes is consistent with previous findings that prasinoviruses exhibit a high degree of conservation in genome structure (Moreau et al. 2010), with the notable exception of McV-KB2, the divergence of which is also reflected in its distinctive G+C content and coding density. Electron micrographs of McV-SA1 established a virion capsid diameter of ∼142-– 160 nm (Fig. 3).

**Figure 2.**
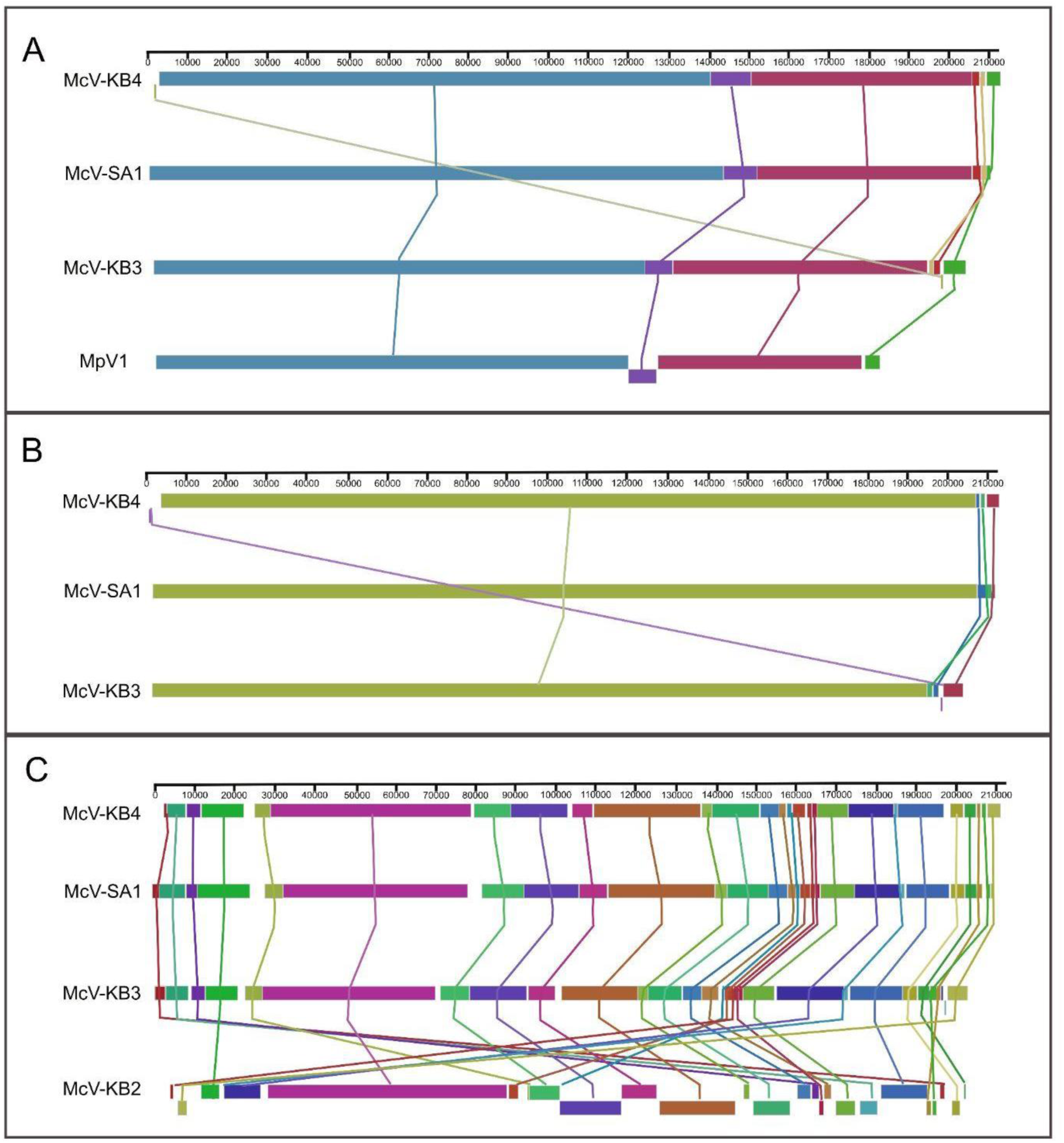
Whole genome alignments created with the progressiveMauve algorithm. Colored blocks represent regions conserved across genomes. (A) An alignment of McV-KB3, McV-KB4, and McV-SA1 with the published genome of *Micromonas pusilla* virus MpV1. (B) An alignment of McV-KB3, McV-KB4, and McV-SA1. (C) An alignment of all four HiMcV assemblies. Note that the inclusion of McV-KB2 generated a greater number of colored blocks because that genome possesses many inversions and rearrangements relative to the other three.

**Figure 3.**
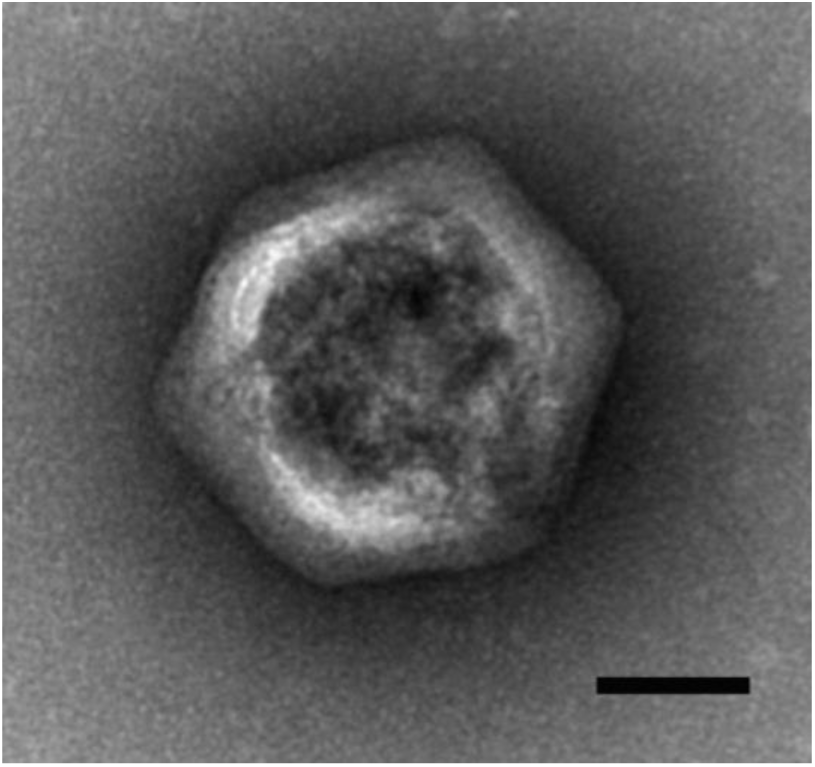
Electron micrograph of McV-SA1 particle. Scale bar = 50 nm.

### Phylogeny

The species tree derived from the concatenation of shared prasinovirus and chlorovirus genes shows that McV-KB3, McV-KB4, and McV-SA1 group into a clade, with McV-KB4 and McV-SA1 most closely related to each other, and the clade of these three HiMcVs lies within a larger clade containing all previously published *Micromonas-* and *Ostreococcus*-infecting virus genomes (Fig. 4). Within this larger clade the *Ostreococcus*-infecting viruses form a monophyletic group, while the *Micromonas*-infecting viruses are paraphyletic, consistent with previous analyses (Bellec et al., 2009; Bachy et al., 2021). McV-KB2 is quite divergent from the other HiMcVs, as its closest relative is the clade of *Bathycoccus*-infecting viruses, although it diverged from that group soon after their common ancestor arose from the last common ancestor of all the prasinoviruses. The divergence of McV-KB2 and its grouping with *Bathycoccus* viruses is consistent with the polB phylogeny, as well as the OrthoFinder/STAG-generated species tree (Supplementary Figures S2 & S3). Although McV-KB2 is relatively divergent from the other HiMcVs, it should be noted that it nonetheless overlaps in host range with each of the other three viruses (Fig. 1). We did not assess whether *Bathycoccus* strains in culture collections can be infected by the HiMcVs, or whether known *Bathycoccus* viruses can infect the *Micromonas commoda* strains from Kāne‘ohe Bay. However, as of this writing, no known prasinovirus infects prasinophytes outside of its isolation host’s genus (see Bachy et al., 2021 and references therein).

**Figure 4.**
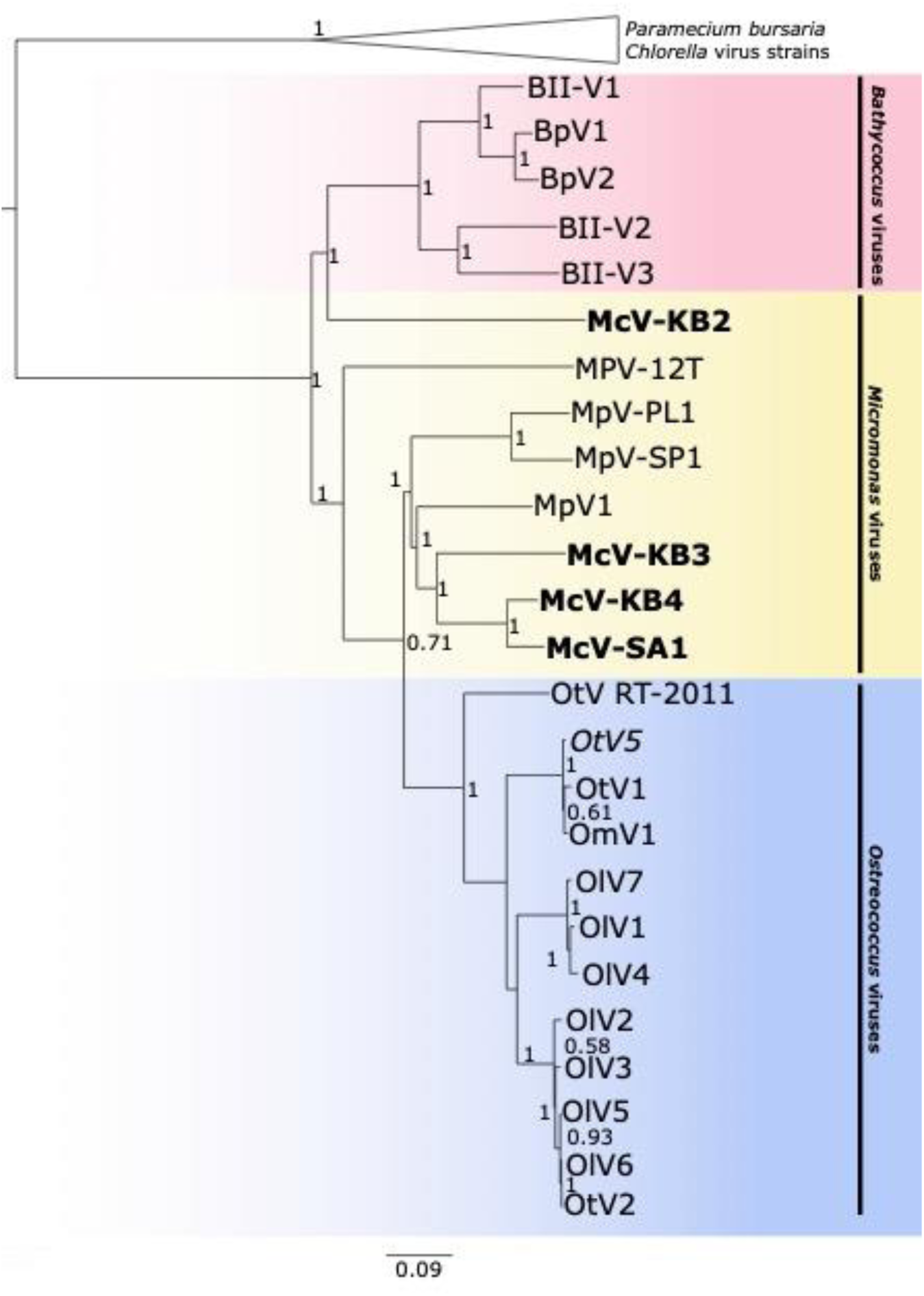
Prasinovirus species tree from concatenated amino acid alignments of the 26 orthogroups shared by all prasinoviruses and chloroviruses in the analysis. Generated with FastTree accessed through Geneious (default settings). Scale bar indicates substitutions per site. The HiMcV isolates are highlighted with bold text.

### HiMcV core genes

Our OrthoFinder analysis found 344 orthogroups that were present within at least one of the HiMcV genomes (Table 2). The four HiMcVs share 152 orthogroups (i.e., the core HiMcV orthogroups), while the other 192 orthogroups are present in three or fewer genomes (Fig. 5). Fifty-six of the HiMcV core orthogroups were found in all prasinoviruses, and 117 were found in all *Micromonas* virus genomes (Table 2). There were 48 orthogroups found in at least one HiMcV that were also found in at least one of the two host genomes, M1 and M2 (Table 2).

**Figure 5.**
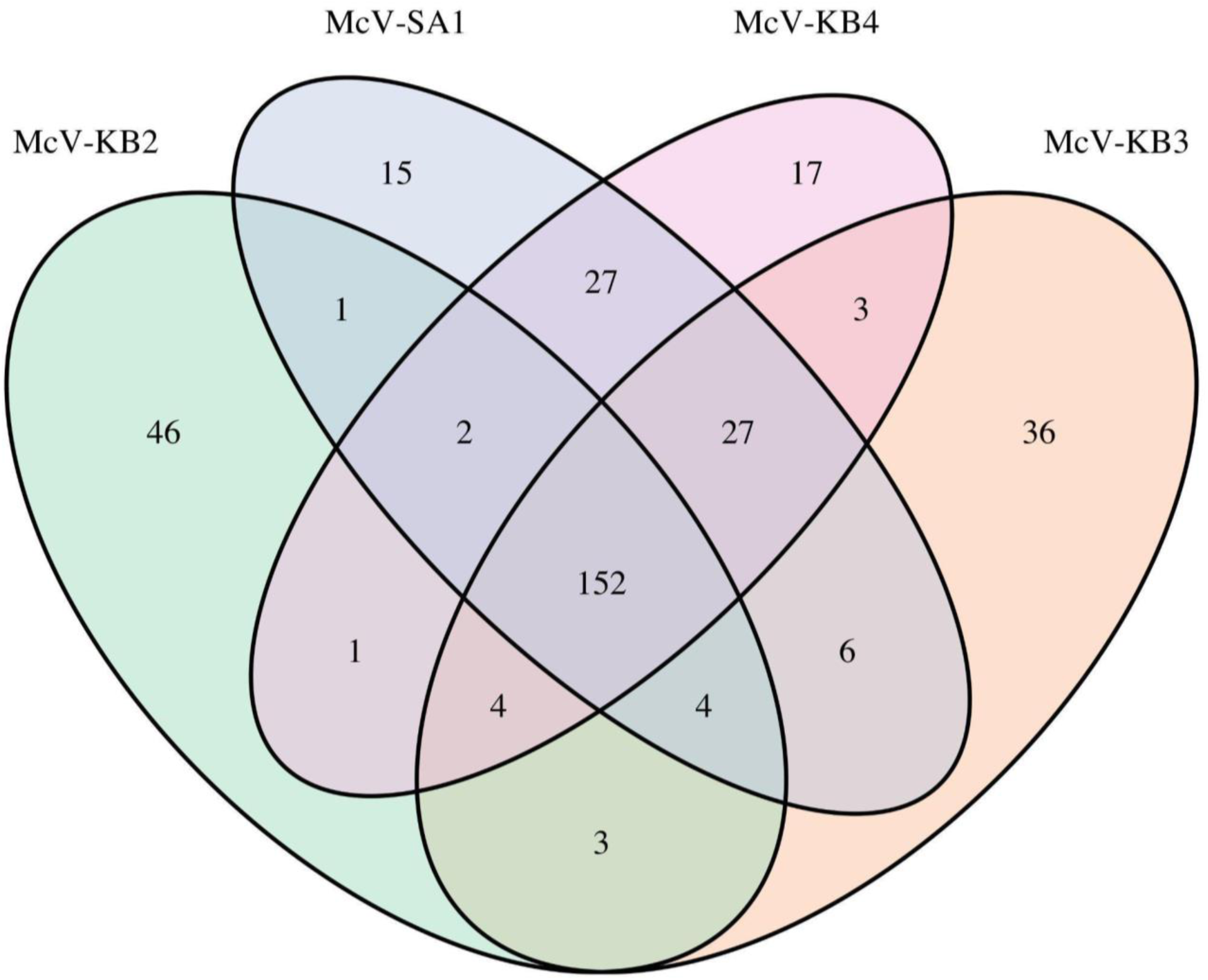
Venn diagram of the number of orthogroups shared by and unique to the four HiMCVs.

**Table 2.** Summary of OrthoFinder results comparing prasinoviruses, chloroviruses, and *Micromonas* hosts M1 and M2. Here the term “core” refers to orthogroups that are present in 100% of genomes within the indicated taxa.

|  | No. of Orthogroups |
| --- | --- |
| Total prasinovirus | 693 |
| Total HiMcV | 344 |
| Core HiMcV | 152 |
| Core *Ostreococcus* viruses | 129 |
| Core *Micromonas* viruses | 117 |
| Core *Bathycoccus* viruses | 99 |
| Core prasinovirus | 56 |
| Core prasinovirus + chlorovirus | 26 |
| HiMcV not found in other prasinoviruses | 61 |
| HiMcV shared with host | 48 |

Among the HiMcVs, McV-KB2 is the most distinct, with 46 unique orthogroups, and McV-KB3 has the second highest number of unique orthogroups (36) which is in concordance with the phylogenetic distances between the HiMcVs (Fig. 4). Host range of the HiMcVs (Fig. 1) did not obviously map onto phylogenetic relatedness or similarity of gene content, perhaps not surprising because of the small number of genomes in our dataset (n=4). McV-KB2 and McV-KB3 both have relatively broad host range, and share five host strains out of the six that each can infect (Fig. 1), but these viruses are distantly related and share only 163 orthogroups (Figs. 4, 5). The two most closely related HiMcVs, McV-KB4 and McV-SA1 (Fig. 4), share 208 orthogroups (Fig. 5) These viruses are similar in having a relatively narrow host range (2 hosts lysed out of 7 total), although they only share 1 host strain out of the 2 that each can infect (Fig. 1).

All 152 core HiMcV orthogroups contain genes that occur in at least one other prasinovirus (Supplementary Table S5). Broadly speaking, core HiMcV orthogroups with functional annotation encompass a diverse suite of viral biology, such as virion structure (major capsid protein), genome replication (DNA polymerase, DNA primase), RNA processing and transcription (mRNA capping enzyme, transcription factors, RNAse H), breakdown of host polymers (nucleases, proteases), nucleotide metabolism (dUTPase, dCMP deaminase, thymidylate synthase, ribonucleotide reductase), carbohydrate metabolism (mannitol dehydrogenase), lipid metabolism (glycerophosphoryl diester phosphodiesterase, phospholipase), and glycosylation (glycogen phosphorylase B, nucleotide-diphospho-sugar transferases). All four HiMcVs possess a cAMP-dependent Kef-type K+ transporter that is found in M1 and M2, as well as *Micromonas* virus strains SP1 and MpV1.This transporter is known in bacteria to help protect cells from electrophilic compounds (Rasmussen, 2023). Potassium channels are the most common type of membrane transporters encoded by viral genomes, with substantial diversity, and likely multiple origins.(Greiner et al., 2018). In *Paramecium bursaria Chlorella* viruses a potassium channel depolarizes the cell membrane during infection, preventing infection of the cell by multiple viruses (Greiner et al., 2009). Thus, the Kef-type potassium transporter may play a significant role in the infection strategy of HiMcVs.

Another noteworthy core HiMcV orthogroup is annotated as a high-light inducible protein in refseq_protein, and as chlorophyll a-b binding protein in InterPro. This orthogroup occurs in M1 and M2 hosts and is present in the assembly for *Micromonas* virus MpV1. Chlorophyll a-b binding proteins are found in the light harvesting complex of photosystem II and play an important role in regulating excitation energy under fluctuating light levels (Liu and Shen, 2004). There is growing evidence that the high-light inducible / chlorophyll a-b binding protein plays an important role in the viruses of eukaryotic phytoplankton, as it has been found in other prasinoviruses, *Chrysochromulina ericina* virus CeV-01B (family Mesomiviridae), and a variety of giant virus metagenome-assembled genomes (Gallot-Lavallée et al., 2017; Moniruzzaman et al., 2020). High-light inducible proteins are known in cyanobacteria to be activated in the presence of excessive photon energy, which suggests that a virus could utilize this protein to protect the cell from photodamage during the viral infection cycle (Dolganov et al., 1995). Additionally, similar proteins in micro- and macro-algae are differentially expressed in nutrient-poor environments (Varsano *et al*., 2006; He *et al*., 2023).

A putative PhoH-like phosphate starvation-inducible protein is also among the HiMcV core sequences. This protein has been seen previously in some, but not all, prasinoviruses (Monier et al., 2012) and is common among marine phages (Goldsmith et al., 2011), potentially enhancing viral infection under low-phosphate conditions, or reflecting the fact that viruses tend to have a higher stoichiometric P content than their hosts (Jover et al., 2014), and therefore benefit from enhanced P supply even if the host is not P-limited. While genes in this family are common in eukaryotic phytoplankton, it appears that our host genomes do not contain orthologs to the PhoH-like sequences found in our HiMcVs. Previously studied prasinovirus versions of this gene appear to be host-derived, which may mean that the HiMcVs obtained it from other *Micromonas* hosts in the environment (Monier et al., 2012) or possibly from other viruses during coinfection, which has been observed for intein acquisition in prasinoviruses (Clerissi *et al*., 2013).

A final notable gene shared among all four HiMcVs is an alternative oxidase. Within this orthogroup, McV-KB2, McV-KB3, and McV-SA1 had one ortholog each, while McV-KB4 had three orthologous sequences, the host strain M1 had one sequence, and M2 had two sequences. Alternative oxidases have been previously found in cyanophages (Puxty et al., 2015), and these enzymes are thought to reduce photodamage to the electron transport chain under stressful conditions, which, in the case of infections, may arise from viral inhibition of photosystem I and ferredoxin NADP+ reductase (FNR) (Wang et al., 2023). Our analysis found alternative oxidases in only one other prasinovirus beyond the HiMcVs, the coastal Mediterranean *Ostreococcus tauri* virus RT-2011.

In the gene tree resulting from our alignment of diverse alternative oxidases, including ubiquinol oxidases (AOX) and plastoquinol terminal oxidases (PTOX), all prasinovirus sequences appear in the same clade (Fig. 6). The closest relatives of this clade are sequences from *Micromonas* isolates, including one of the sequences from M2. This group, containing prasinovirus and *Micromonas* PTOX sequences, is a sister clade to one containing *Bathycoccus* and *Ostreococcus* sequences, although support for the node joining these clades is low (0.21). Within the prasinovirus clade there are three divergent sequences from McV-KB4, suggesting several duplication events.

**Figure 6.**
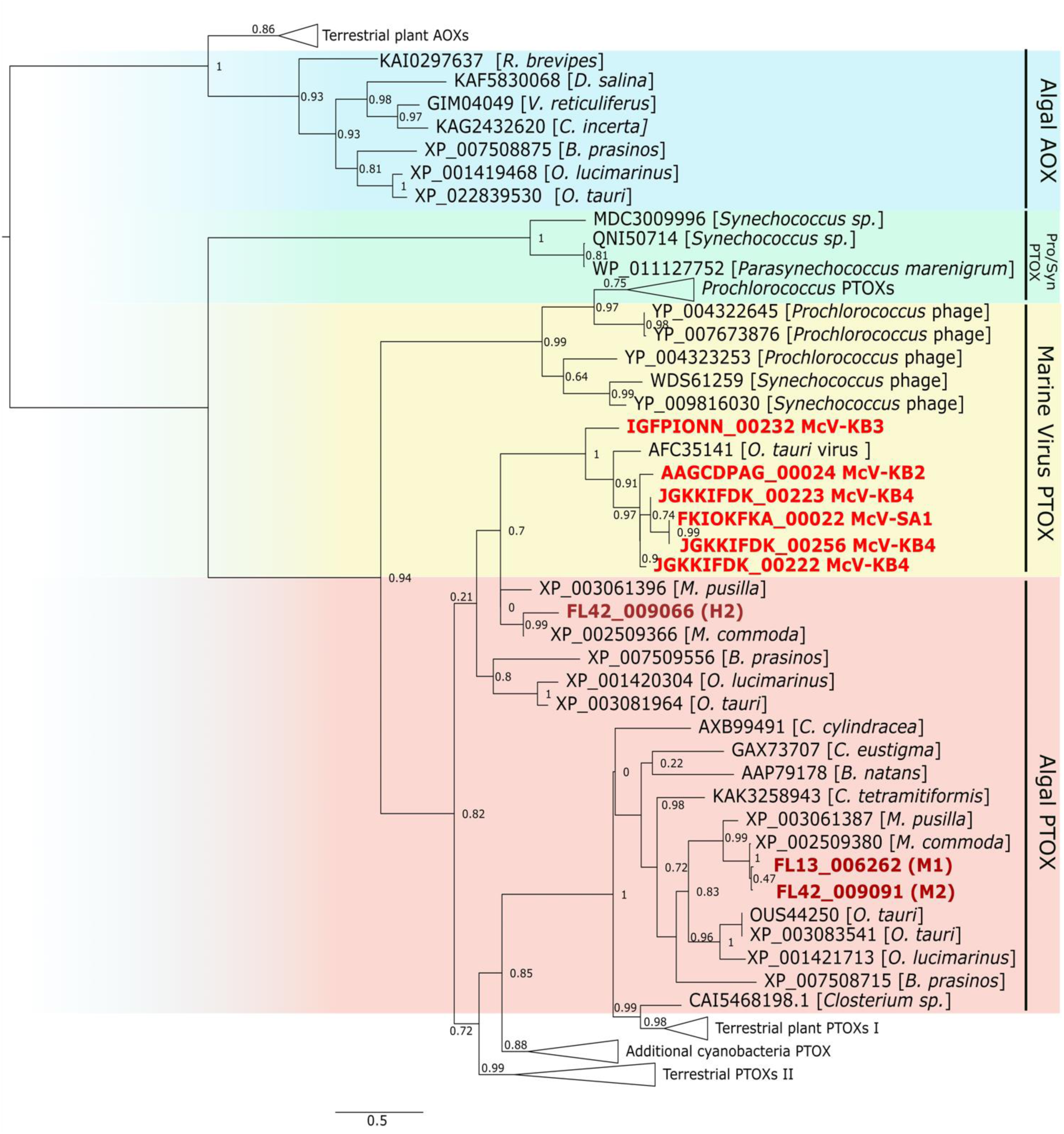
Alternative oxidase/plastoquinol terminal oxidase gene tree, including mitochondrial ubiquinol oxidase (AOX) from plants and eukaryotic phytoplankton, and plastoquinol terminal oxidase (PTOX) from plants, eukaryotic phytoplankton, cyanobacteria, prasinoviruses, and cyanophages. Vertical labels to the right of the tree are used to aid in visual interpretation and do not necessarily indicate a close relationship among genes. Gene sequences were aligned in MAFFT and trimmed with GoAlign, and tree created in FastTree. Node support values reflect FastTree Shimodaira-Hasegawa test values. UHM strains are shown in bold, with sequences form virus strains in red and host strains in maroon. The tree was rooted using the split between AOX and PTOX sequences. Scale bar indicates substitutions per site.

Further inspection of the AOX/PTOX phylogeny reveals several important features. First, there is a monophyletic clade containing AOX from prasinophytes, other algae, and plants, consistent with an ancient divergence between AOX and PTOX (Nobre et al., 2016). Second, within the PTOX clade there is a relatively early divergence by *Synechococcus/Parasynechococcus* sequences, placing them in a different part of the tree from the *Synechococcus* phage sequences, which are more closely related to *Prochlorococcus* phage and *Prochlorococcus* cellular sequences, which was also noted in a prior analysis (Puxty et al., 2015). Third, after the divergence of *Synechococcus* sequences there is a split between a clade containing the *Prochlorococcus* and cyanophage sequences and a clade containing eukaryote and prasinovirus PTOX sequences as well as other cyanobacterial sequences. Finally, the eukaryote+prasinovirus+cyanobacteria clade contains two main branches, and the two branches possess different paralogs of PTOX from isolates of *Micromonas* (including M2), *Ostreococcus*, and *Bathycoccus*. This split may represent an ancient duplication event resulting in two PTOX copies, one of which was acquired by prasinoviruses. Based on the structure of the phylogeny it may be the case that prasinovirus and cyanophage PTOX genes were acquired through distinct gene transfer events.

### Notable non-core and unique HiMcV genes

There are 61 non-core HiMcV orthogroups that represent proteins not previously found in prasinovirus isolates (verified through a secondary nr BLAST search), which we will refer to as “unique”. Limited functional information is available for the 61 unique orthogroups, as 40 have no BLAST hits against refseq_protein, eight have BLAST hits to hypothetical proteins, and the remaining 13 include many with low-identity (< 30% amino acid identity) BLAST hits. Results from the InterPro databases are comparable, although many of the proteins are predicted to be membrane-bound. One notable unique orthogroup is a phosphate:sodium symporter in McV-SA1 that is also found in both M1 and M2 host genomes. Top BLAST hits for this McV-SA1 CDS in both nr and refseq_protein are from previously published strains of *Micromonas*, including RCC299, a pelagic strain from the equatorial Pacific, and CCMP1445, a coastal strain from the North Atlantic. This CDS has no orthologs among the other prasinovirus genomes, which may indicate that the McV-SA1 symporter gene has a recent cellular origin via horizontal gene transfer. It should be noted that our data set includes a second orthogroup with prasinovirus phosphate transporter sequences that appear distinct from the McV-SA1 symporter, as an alignment of the two orthogroups indicated low sequence identity, albeit with scattered matching residues (results not shown). This second orthogroup includes phosphate transporter sequences from BpV1 (HM004431), BII-V1 (MK522034), OlV6 (HQ633059), OlV5 (NC_020851), OtV2 (FN600414), OlV4 (JF974316), OlV2 (NC_028091), and OlV1 (MK514405), as well as sequences from M1 and M2. This orthogroup corresponds to the prasinovirus phosphate transporters from the PHO4 superfamily identified by Monier et al. (2012) in a comparison of phytoplankton virus phosphate transporters. The PHO4 transporters correspond to InterPro family IPR001204, whereas the novel McV-SA1 transporter corresponds to InterPro family IPR003841. Therefore, the McV-SA1 phosphate:sodium symporter likely indicates a separate acquisition of a phosphate transporter by prasinoviruses, potentially with different uptake affinity or other physiological differences.

Other unique HiMcV orthogroups with predicted functions include a putative bax inhibitor-1 (McV-KB2), N-6 DNA methylase (McV-KB2), polyamine aminopropyltransferase (McV-KB3), adenosylmethionine decarboxylase (McV-KB3), and a glycosyltransferase (McV-SA1). Bax inhibitor-1 is a conserved inhibitor of programmed cell death, and viruses such as deerpox (Banadyga et al., 2011) and cytomegalovirus (Ma et al., 2012) are known to encode other proteins that suppress bax, thereby countering the elimination of infected cells by apoptosis. Top BLAST hits to the McV-KB2 bax inhibitor-1 gene are sequences from fungi, but the amino acid identity is low (30-35%), making it unclear where McV-KB2 may have acquired the gene from.

The putative adenosylmethionine decarboxylase and polyamine aminopropyltransferase encoded by McV-KB3 may catalyze linked steps in the synthesis of spermidine from putrescine. Spermidine is a polyamine required for cell growth as well as the replication of many viruses. Enzymes related to spermidine synthesis have been found in a variety of phages and eukaryotic viruses (Li et al., 2023), including the chlorovirus PBCV-1 (Baumann et al., 2007), but the McV-KB3 genes appear to be the first reported occurrence in a prasinovirus isolate. The two McV-KB3 genes have BLAST hits to genes from various bacteria and archaea, although the relatively low amino acid identity (∼30-40%) provides little information about the proximate origin of the genes. The other unique orthogroups with putative functions (N-6 DNA methylase and a glycosyltransferase) represent categories of enzymes that are commonly encoded by prasinoviruses, although the uniqueness of these orthogroups indicates these specific genes are not closely related to previously documented prasinovirus genes. A final notable unique orthogroup is a gene found only in McV-KB4 that is orthologous to cyanophage genes of unknown function. This putative CDS has not been seen in other prasinoviruses, and the closest database hit is a hypothetical protein from *Prochlorococcus* phage strain P-SSM2, which infects low-light *Prochlorococcus* ecotypes. The orthogroup may be evidence of gene exchange between cyanophage and a eukaryotic virus.

While none of the unique orthogroups have been previously seen in isolates, most (42 of 61) have homologs in the GOEV database. Eight of these genes belong to McV-KB2, ten to McV-KB3, and one to McV-SA1. Seventeen of these genes have limited information from InterPro databases, e.g., indicating a membrane-bound protein and/or a coil. Of the forty-two unique orthogroups with GOEV hits, thirty-five were homologous to MAG-derived GOEV gene sequences classified as belonging to the genus Prasinovirus, while the remaining fell into various taxa in the phylum *Nucleocytoviricota*, with two sequences from Imitevirales.

### HiMcV orthogroups shared with their host genomes

A total of 48 HiMcV orthogroups contain genes also found in the M1 and/or M2 host genomes, 27 of which are shared among all 4 HiMcVs. In total M1 shared 27 orthogroups with all four HiMcV strains, with 19 additional orthogroups shared between M1 and at least one other virus (Supplementary Figure S4). Results from comparison with M2 were similar, with 24 orthogroups shared between M2 and all four viruses, and with 22 orthogroups shared between M2 and at least one virus (Supplementary Figure S5). The number of shared orthologs is comparable to results from Moreau et al. (2010), in which *Micromonas pusilla* virus MpV1 shared 56 CDS with *Micromonas* sp. strain RCC1109. Forty-five of the HiMcV-host shared orthogroups contain genes found in previously published prasinoviruses, as made apparent by refseq_protein and nr BLAST hits. Some of these orthogroups were described in the section ‘HiMcV core genes’ (chlorophyll a-b binding protein, PTOX, cAMP-dependent Kef-type K+ transporter). In general, the orthogroups shared with hosts are associated with a variety of cellular processes such as protein modification/regulation/processing (N-myristoyltransferase, ubiquitin, cysteine protease, ATP-dependent metalloprotease FtsH), glycosylation (nucleotide-sugar epimerases and transferases), amino acid synthesis (dehydroquinate synthase), nucleotide metabolism (dCMP deaminase, thymidine kinase), transcription regulation (transcription factors), stress response (heat shock protein 70, rhodanese, superoxide dismutase, mannitol dehydrogenase), nucleic acid processing (exonuclease, ribonucleotide reductase, DNA polymerase family X), photosynthesis (PTOX, chlorophyll a/b binding protein), and perhaps countering host defenses (methyltransferases) (Supplementary Table S7).

There are three HiMcV orthogroups not found in other prasinoviruses that are found in both hosts, which include the aforementioned phosphate:sodium symporter found in McV-SA1, as well as two orthogroups shared with McV-KB2. One of the McV-KB2:host orthogroups contains only hypothetical protein sequences with no hits in NCBI or InterPro databases. The other McV-KB2:host orthogroup contains sequences that are annotated as chlorophyllide a oxygenase (CAO) for M1 and M2. CAO converts chlorophyll a to chlorophyll b, an important accessory pigment in green algae (Jeffrey et al., 2011), which may mean that CAO supports light adsorption during infection by McV-KB2. However, amino acid identity between the McV-KB2 and host sequences is 17.89%, suggesting these sequences may not be truly orthologous. Local collinearity throughout the alignment of these sequences may explain their ultimate clustering into an orthogroup. If McV-KB2 indeed encodes for CAO it would be the first virus reported to have this gene.

### Genes differentiating prasinoviruses that infect different host genera

In total there were 693 orthogroups in our analysis that occurred in at least one prasinovirus. Linear models relating orthogroup representation for each prasinovirus to host genus found 170 orthogroups that differed significantly between viruses of the three host genera (p < 0.05; Supplementary Table S8). Therefore, 25% of prasinovirus orthogroups were significantly associated with host genus identity, suggesting that a substantial portion of the genome is involved in adapting to infect different host genera in the same taxonomic order. Twenty-eight orthogroups that differ strongly between host genera (p < 0.001) and that also have functional annotations (Supplementary Table S3) exemplify the diversity of functions that relate viral gene content to host identity. For example, orthogroups that are absent in *Bathycoccus* viruses but present in most or all *Micromonas* and *Ostreococcus* viruses include asparagine synthetase (nitrogen and amino acid metabolism), dCMP deaminase (nucleotide metabolism), DNA polymerase X (potentially for base excision repair; Fernández-García et al., 2017), nucleotide-diphospho-sugar transferases (glycosylation), RNAse H (RNA processing), NTP pyrophosphohydrolase (potentially involved in stress response), and a protein with a rhodanese-like domain (potentially involved in stress response). Orthogroups that are present in most or all *Micromonas* viruses but absent/rare in *Ostreococcus* and *Bathycoccus* viruses include a glycerophosphodiester phosphodiesterase (lipid metabolism), mannitol dehydrogenase (potentially involved in stress response), ubiquitin, a protein with a zinc finger C2H2-type domain (potential transcription factor), a protein with an integrin alpha domain (potentially used for attachment to the host), and a putative tail fiber protein (potentially used for attachment to the host). Therefore, it may be the case that many stages of the viral life cycle, such as attachment to host receptors and manipulation of host metabolism and defenses, are involved in (co)evolution to infect different host genera.

We used unsupervised clustering analysis to further understand how gene content varies among the prasinovirus genomes that we analyzed (Fig. 7). Consistent with the many differences we found between viruses infecting host genera (Supplementary Table S3), unsupervised clustering largely groups viruses by host genus, with the exception of one *Ostreococcus tauri* virus that occurs in the *Micromonas* virus cluster. In our phylogenetic analysis this strain, *Ostreococcus tauri* virus RT-2011 (JN225873.1), is relatively divergent from the clade containing the other 12 *Ostreococcus* viruses (Fig. 4). It is possible that OtV RT-2011 retained gene content similar to *Micromonas* viruses while the other *Ostreococcus* viruses evolved more *Ostreococcus*-specific gene content. Finally, the clustering results again emphasize the uniqueness of McV-KB2 relative to the other HiMcVs, as the other three HiMcVs cluster together, while McV-KB2 is grouped with the genome of MpV-12T, a strain isolated from the coast of the Netherlands.

**Figure 7.**
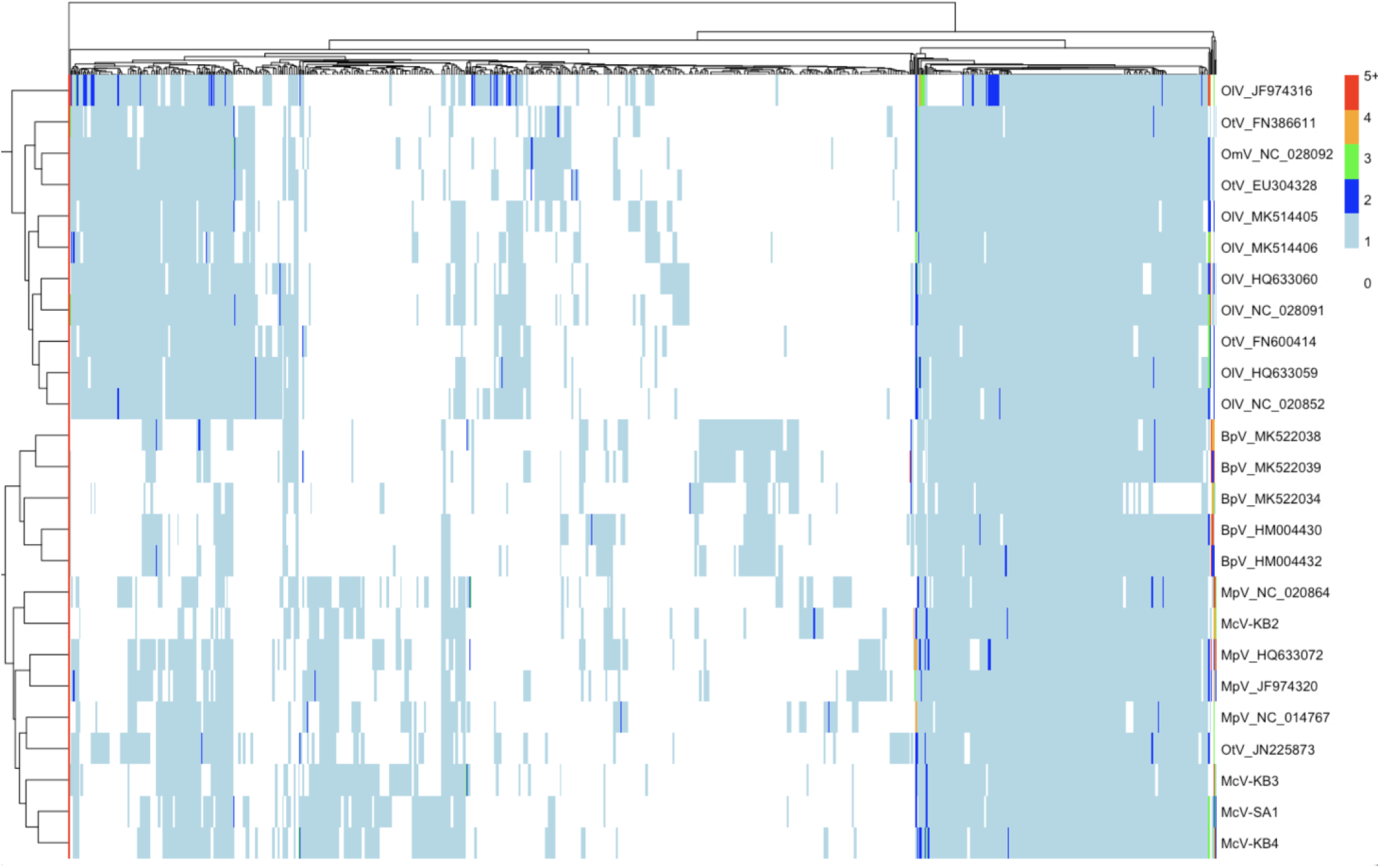
Clustered heatmap of 693 prasinovirus orthogroups. Colors in the heatmap represent the number of gene copies in each orthogroup per genome, with warmer colors being higher. The dendrogram on the left-hand side reflects similarity of virus strains based on orthogroup composition, and the dendrogram at the top clusters orthogroups by similarity in patterns of occurrence across strains.

### Distribution of HiMcV sequences in the world ocean

Using CoverM we observed 115 instances of metagenomic reads mapping to our HiMcVs from the combined Station ALOHA, GEOTRACES, and Tara Oceans datasets (Supplementary Table S9). Although our viruses were isolated at, or relatively near, Station ALOHA, only 8 out of 185 samples at this location included hits to a HiMcV (Fig. 8). The lower relative occurrence of HiMcVs in the Pacific, compared to the Atlantic, was further suggested by no hits from samples in the GEOTRACES transect GP13 in the South Pacific or any of the Pacific Tara Oceans stations (Fig. 8).

**Figure 8.**
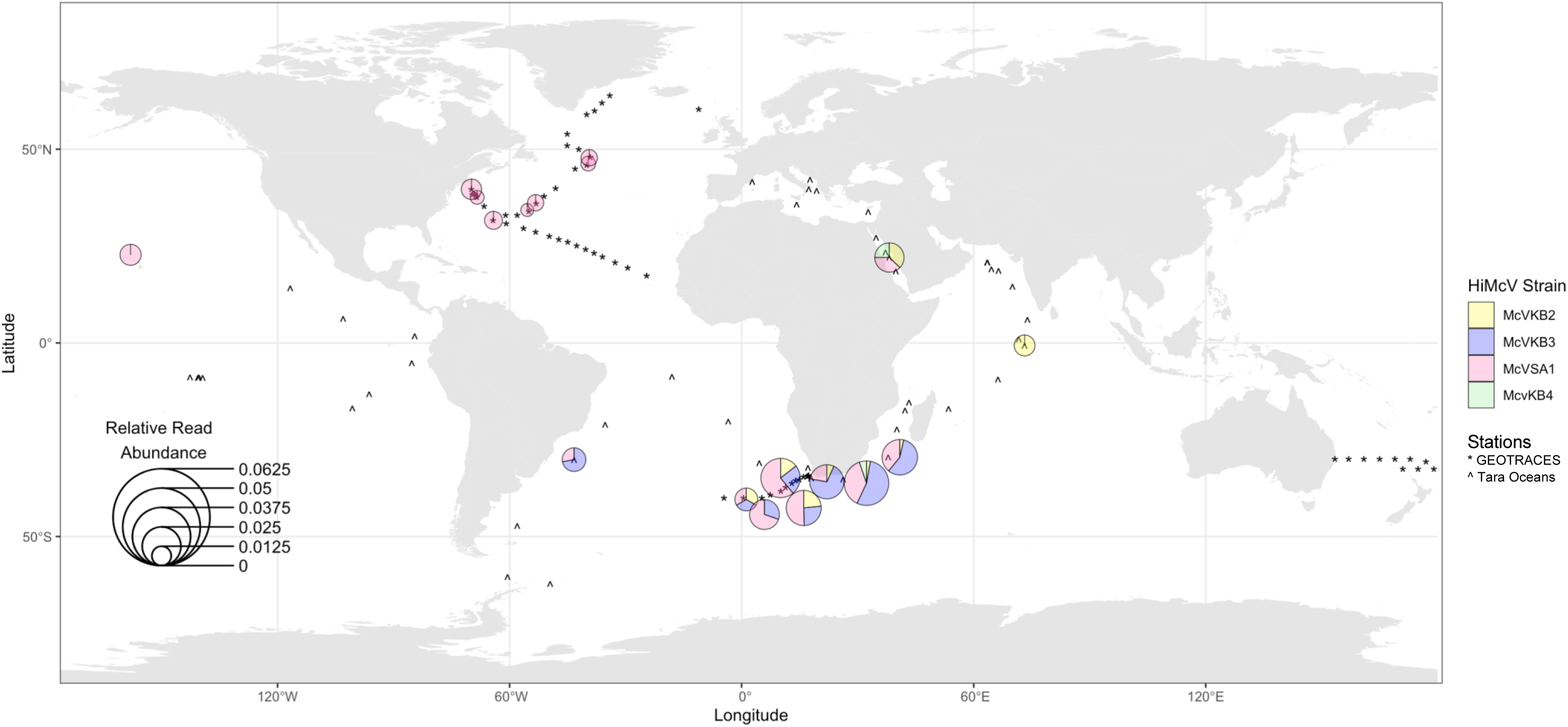
Distribution of HiMcV strains in metagenomic samples from GEOTRACES, Tara Oceans, and Station ALOHA. Pie charts represent strain composition at each location, based on mean relative abundance of each strain. The size of each pie chart indicates the summed relative abundance of HiMcV transcripts. For HOT data, relative abundance was averaged over 8 samples from different time points. Stations where no HiMcV strains were detected are represented with an asterisk for GEOTRACES and a carrot for Tara Oceans. The smaller pie charts in the North Atlantic and Station ALOHA all contain reads only from McV-SA1. Pie chart locations are approximate, as coordinates were adjusted to prevent overlap.

The HiMcV strain isolated from the open ocean, McV-SA1, accounts for over half of all HiMcV hits in the metagenomic datasets. This strain was also the only HiMcV found in the North Atlantic and North Pacific basins. Interestingly, McV-KB4, which is sister to McV-SA1 in the phylogeny (Fig. 4), is underrepresented, with only three hits. This may be evidence that genome similarity does not predict similar ecology. HiMcV strain diversity and relative abundance tended to increase on the edges of subtropical gyres, in transition zones, or near islands (Fig. 8), which tend to be more productive compared to the pelagic ocean.

In addition to our CoverM search of metagenomes, we compared McV isolate genomes to entire GVMAG assemblies in the GOEV database and found one GVMAG, TARA_AON_NCLDV_00048, that is closely related to McV-SA1. All contigs of this GVMAG could be mapped to the McV-SA1 genome, resulting in 71% coverage with 92% nucleotide identity (Supplementary Fig. S6). Ha et al. (2023) found that this GVMAG was the 8th most common *Algavirales* virus and the 15th most common *Nucleocytoviricota* virus in the bioGEOTRACES metagenome survey, out of 696 GVMAGs and 1382 *Nucleocytoviricota* isolate genomes to which metagenome reads were mapped. The TARA_AON_NCLDV_00048 genome occurred in 42 out of 480 samples. For comparison, 76 of our CoverM hits came from the bioGEOTRACES database, 47 of which came from McV-SA1. Similar to McV-SA1, the *Tara* GVMAG occurred in the three Atlantic bioGEOTRACES transects, but not the sole Pacific transect.

## Discussion

Our study highlights the genomic diversity of prasinoviruses in several ways. Novel viruses from the same location, with overlapping host range, exhibited substantial variation in gene content, and one of the newly sequenced genomes (McV-KB2) was relatively divergent from all previously sequenced prasinoviruses. The 61 HiMcV orthogroups that are novel to prasinoviruses also indicate that substantial functional diversity remains to be discovered in this clade, and judging from the novel orthogroups with functional annotations (a phosphate transporter, an apoptosis inhibitor, enzymes from spermidine synthesis) the prasinovirus pangenome likely includes diverse mechanisms of virocell manipulation. The majority of the 61 novel orthogroups had homologs in metagenome-assembled genomes from the ocean, further suggesting a high level of uncultivated genetic diversity. Finally, the various orthogroups that differ among viruses infecting *Micromonas*, *Ostreococcus*, and *Bathycoccus* point towards a better understanding of the functional basis of viral diversification. Viral gene content is strongly tied to host genus identity, and orthogroups unique to each genus suggest these may have diverse functions, such as virion attachment, manipulating host stress responses, and metabolism of components needed for virion construction.

Although the sample size is relatively small, it is noteworthy that all HiMcVs carry one or more orthologs to a host PTOX, and all HiMcVs also encode a high-light inducible chlorophyll a-b binding protein. Among prasinoviruses a PTOX sequence was previously found in only one other isolate (OtV-RT2011), but PTOX is commonly found in cyanophages (Puxty et al., 2015). The common occurrence of this gene in disparate virus clades may indicate an increased need for protection against photodamage for infected photosynthesizers at low latitudes. The occurrence of the high-light inducible chlorophyll a-b binding protein present in all HiMcVs may also contribute to maintaining host metabolism under light stress, although at this point it is unknown whether the viral proteins fulfill their annotated functions in infected cells.

Metagenomic data provide evidence that one HiMcV strain, McV-SA1, is relatively common in the global ocean. In the bioGEOTRACES transects this virus occurs at a similar frequency to a closely related uncultivated virus, which was shown to be one of the most common *Nucleocytoviricota* viruses in the those transects (Ha et al., 2023). This suggests that *Micromonas*-infecting viruses are among the most common *Nucleocytoviricota* in the ocean. At the same time, the other three HiMcVs are relatively rare in major ocean basins. Several factors may contribute to the relatively small number of hits against the utilized datasets. These three isolates (McV-KB2, McV-KB3, and McV-KB4) were isolated from a coastal site and may be more abundant in coastal locations compared to the primarily open ocean metagenome stations. The relatively productive region near South Africa, surveyed in the bioGEOTRACES GA10 transect, contained the highest abundance and diversity of HiMcVs, consistent with *Micromonas* being more common under nutrient-rich conditions (Not et al., 2004; Lopes Dos Santos et al., 2017). Therefore, further surveys in productive waters may capture more sequences that map to our HiMcVs.

In conclusion, our study shows that isolating and sequencing new viruses even within a relatively well-studied clade (the prasinoviruses) improves our knowledge of marine viral gene content and genome evolution. Such isolates also provide resources for future functional genomic studies that can resolve questions about putative gene functions and experimental studies to better understand virus contributions to phytoplankton ecology.

## Acknowledgments

This work was supported by NSF grants OCE 1559356 (to GFS and KFE), OCE 2129697 (to KFE, GFS, and CRS), RII Track-2 FEC 1736030 (to GFS, KFE, and SWP) and a Simons Foundation Investigator Award in Marine Microbial Ecology and Evolution (to KFE). Special thanks to M. Marston, R. Chong, and K. Selph for comments on previous versions of this manuscript. Thank you to Brewster Kingham and the staff of the University of Delaware Sequencing and Genotyping Center (RRID:SCR_012230) for sequencing our *Micromonas* cell lines. Support from the University of Delaware Bioinformatics Data Science Core Facility (RRID:SCR_017696) including access to additional computational resources was made possible by Delaware INBRE (NIH/NIGMS P20GM103446), the State of Delaware, and the Delaware Biotechnology Institute.

## Supplementary Materials

**Supplementary Table S1.** Antibiotic recipe used to clean *Micromonas* culture of bacteria and associated phage. This recipe was developed by colleagues at Observatoire océanologique de Banyuls-sur-Mer.

| Antibiotic name | Final concentration<br>(µg/mL) | Mass (g) in 1000X<br>stock (10mL) |
| --- | --- | --- |
| Ampicillin | 50 | 0.5 |
| Gentamicin | 50 | 0.5 |
| Kanamycin | 20 | 0.2 |
| Neomycin | 100 | 1 |

**Supplementary Table S2.** Strain information for prasinovirus and chlorovirus strains used in OrthoFinder and phylogenetic analysis.

| Accession | Strain Name | Strain<br>Abbreviation | Host Genus | Authors |
| --- | --- | --- | --- | --- |
| HM004430 | *Bathycoccus* sp. RCC1105 virus | BpV2 | *Bathycoccus* | Moreau et al. (2010) |
| HM004432 | *Bathycoccus* sp. RCC1105 virus | BpV1 | *Bathycoccus* | Moreau et al. (2010) |
| MK522034 | *Bathycoccus* sp. RCC716 virus 1 | BII-V1 | *Bathycoccus* | Bachy et al. (2019) |
| MK522038 | *Bathycoccus* sp. RCC716 virus 2 | BII-V2 | *Bathycoccus* | Bachy et al. (2019) |
| MK522039 | *Bathycoccus* sp. RCC716 virus 3 | BII-V3 | *Bathycoccus* | Bachy et al. (2019) |
| HQ633072 | *Micromonas pusilla* virus PL1 | MpV-PL1 | *Micromonas* | Henn et al. (2010) |
| JF974320 | *Micromonas pusilla* virus SP1 | MpV-SP1 | *Micromonas* | Henn et al. (2010) |
| NC_014767 | *Micromonas* sp. RCC1109 virus | MpV1 | *Micromonas* | Moreau et al. (2010) |
| NC_020864 | *Micromonas pusilla* virus 12T | MpV-12T | *Micromonas* | Henn et al. (2010) |
| HQ633059 | *Ostreococcus lucimarinus* virus 6 | OIV6 | *Ostreococcus* | Henn et al. (2010) |
| HQ633060 | *Ostreococcus lucimarinus* virus 3 | OIV3 | *Ostreococcus* | Henn et al. (2010) |
| JF974316 | *Ostreococcus lucimarinus* virus 4 | OIV4 | *Ostreococcus* | Henn et al. (2010) |
| MK514405 | *Ostreococcus lucimarinus* virus 1 | OIV1 | *Ostreococcus* | Zimmerman et al. (2019) |
| MK514406 | *Ostreococcus lucimarinus* virus 7 | OIV7 | *Ostreococcus* | Zimmerman et al. (2019) |
| NC_020852 | *Ostreococcus lucimarinus* virus 5 | OIV5 | *Ostreococcus* | Henn et al. (2010) |
| NC_028091 | *Ostreococcus lucimarinus* virus 2 | OIV2 | *Ostreococcus* | Derelle et al. (2015) |
| NC_028092 | *Ostreococcus mediterraneus* virus 1 | OmV1 | *Ostreococcus* | Derelle et al. (2015) |
| EU304328 | *Ostreococcus tauri* virus 5 | OtV5 | *Ostreococcus* | Derelle et al. (2015) |
| FN386611 | *Ostreococcus tauri* virus 1 | OtV1 | *Ostreococcus* | Weynberg et al. (2009) |
| FN600414 | *Ostreococcus tauri* virus 2 | OtV2 | *Ostreococcus* | Weynberg et al. (2009) |
| JN225873 | *Ostreococcus tauri* virus RT-2011 | OtV-RT2011 | *Ostreococcus* | Thomas et al. (2011) |
| DQ491002 | *Paramecium bursaria* *Chlorella* virus NY2A | NY2A | *Paramecium bursaria*<br>*Chlorella* | Van Etten et al. (2006) |
| DQ491003 | *Paramecium bursaria* *Chlorella* virus<br>AR158 | AR158 | *Paramecium bursaria*<br>*Chlorella* | Van Etten et al. (2006) |
| DQ890022 | *Paramecium bursaria* *Chlorella* virus FR483 | FR483 | *Paramecium bursaria*<br>*Chlorella* | Fitzgerald et al. (2007) |
| NC_000852 | *Paramecium bursaria* *Chlorella* virus 1 | PbCV1 | *Paramecium bursaria*<br>*Chlorella* | Yanai-Balser et al. (2010) |

**Supplementary Table S3.**
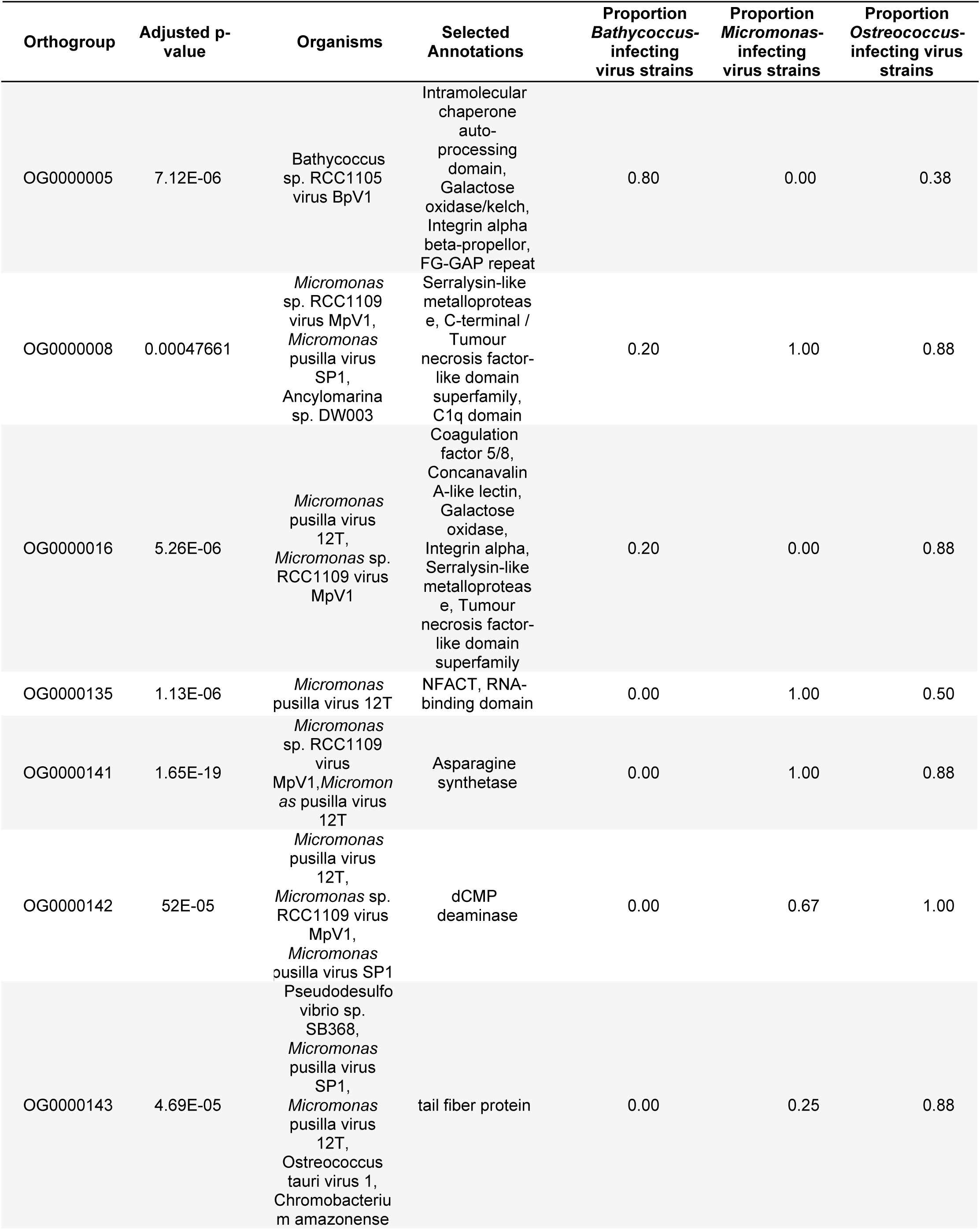
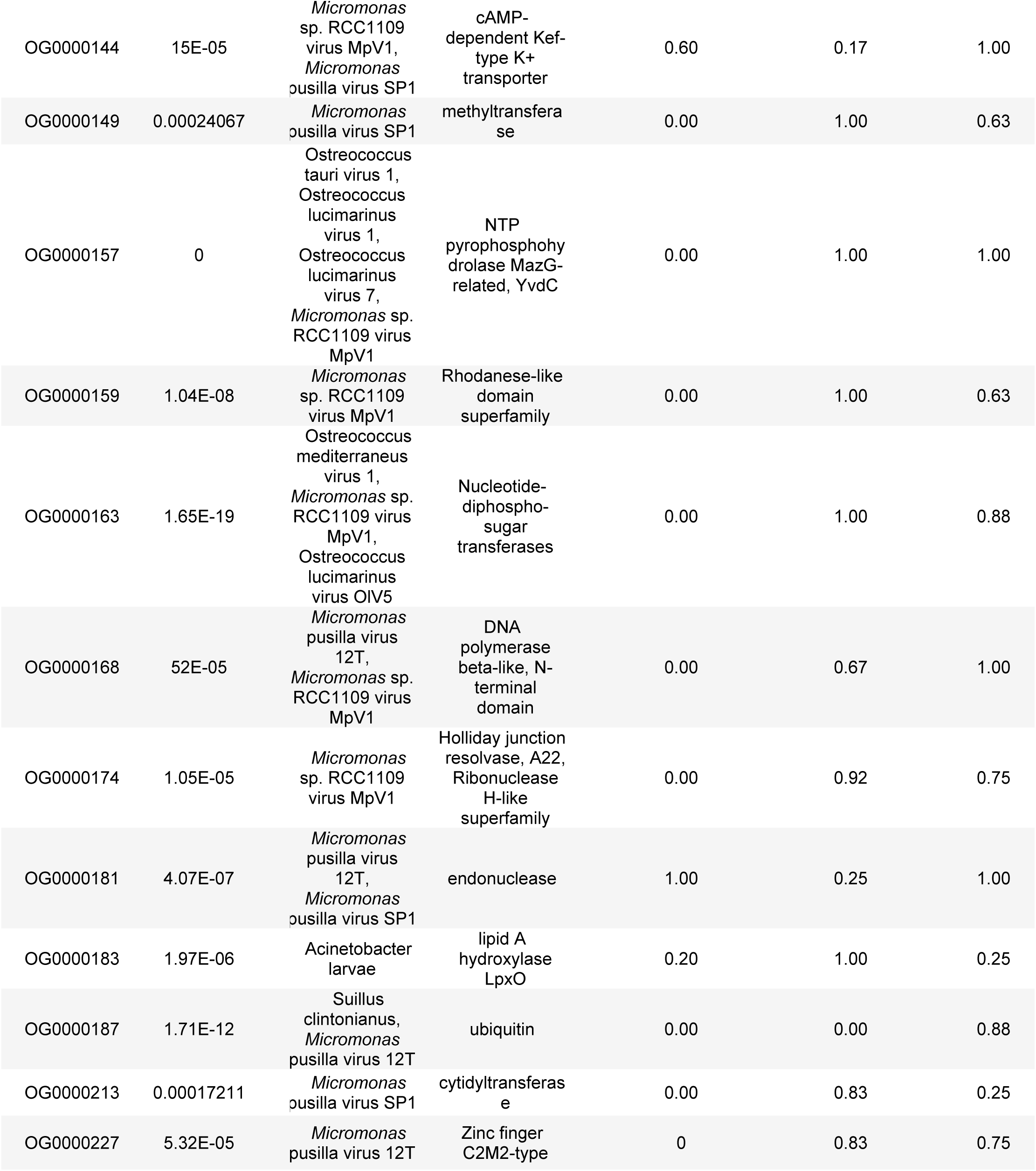
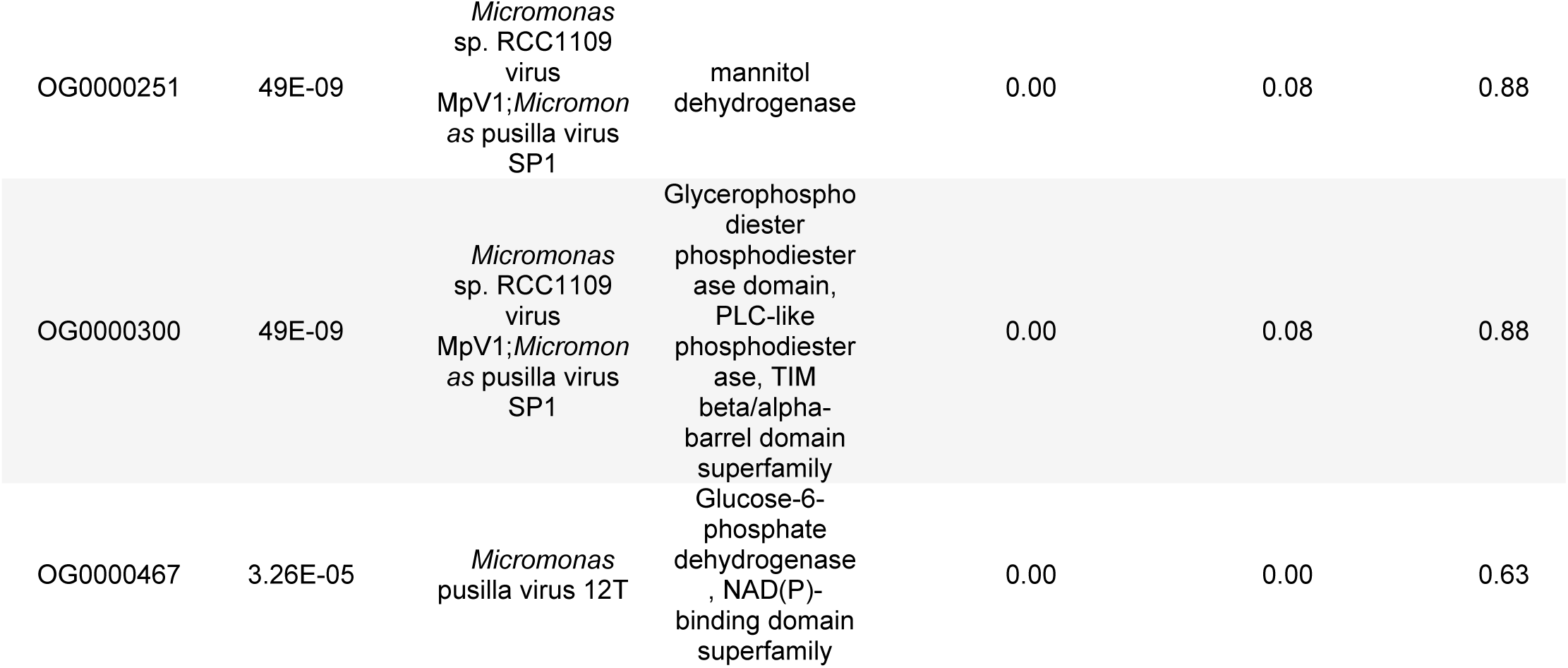
Orthogroups that exhibit highly significant differences between viruses infecting different host genera (p < 0.001), and which possess putative functional annotations. Reported for each orthogroup is the p-value (adjusted for false discovery rate) from a chi-square likelihood ratio test comparing occurrence across host genera, the organism(s) associated with the top refseq_protein database hits, selected annotations from refseq_protein and InterPro, and the proportion of strains infecting a specific host genus that have sequences present in the orthogroup.

**Supplementary Table S4.** Full metagenomic dataset searched with CoverM, including Sequence Read Archive (HOT and GEOTRACES) and European Read Archive (Tara Oceans) accession numbers and metadata. The spreadsheet can be found online here and through the attached csv file.

**Supplementary Table S5.** Orthogroups found in all four HiMcVs (i.e., core HiMcV orthogroups). Table includes top hits from refseq_protein BLAST, information from InterPro member databases, and HiMcV putative gene IDs. The spreadsheet can be found <u>here</u> and through the attached csv file.

**Supplementary Table S6.** Orthogroups not shared by all four HiMcVs (i.e., non-core HiMcV orthogroups), including those not found in other prasinoviruses. Table includes top hits from refseq_protein BLAST, GOEV, information from InterPro member databases, and HiMcV putative gene IDs. Unique orthogroups are highlighted in yellow. The spreadsheet can be found <u>here</u> and through the attached csv file.

**Supplementary Table S7.** Orthogroups shared between HiMcVs and *Micromonas* hosts M1 and M2 Table includes top hits for HiMcVs from refseq_protein BLAST, information from InterPro member databases, HiMcV putative gene IDs, as well as the numbers of host and HiMcV strains with sequences present in each orthogroup. The spreadsheet can be found <u>here</u> and through the attached csv file.

**Supplementary Table S8.** Comparison of orthogroup occurrence across viruses infecting different host genera. Columns for each prasinovirus strain contain sequence count data for each orthogroup (i.e., values > 1 indicate multiple paralogs per strain). Raw p-values, p-values adjusted for false discovery rates, and sequence annotation are included. The corresponding spreadsheet can be found here and through the attached csv file.

**Supplementary Table S9.** Metagenome samples containing reads that mapped successfully to HiMcV assemblies, using the CoverM criteria of 95% nucleotide identity and 20% cover. Table includes strain name of virus, GenBank NCBI SRA or European Read Archive accessions, percent of unmapped reads from each run, relative abundance of reads mapping to HiMcV assembly, and the name of the metagenomic data set. The BIOGEOTRACES (BGT) dataset are from Biller et al. (2018),the Hawaii Ocean Timeseries (HOT) data set is from Mende et al. (2017), and the Tara data set is from Brum et al. (2015).

| Genome | SRA/ENA_Accession | Unmapped(%) | RelativeAbundance(%) | DataSet |
| --- | --- | --- | --- | --- |
| McVKB2 | SRR5788033 | 99.9834 | 0.003044991 | BGT |
| **McVKB2** | SRR5788075 | 99.98971 | 0.001268915 | BGT |
| **McVKB2** | SRR5788032 | 99.984024 | 0.002611603 | BGT |
| **McVKB2** | SRR5788130 | 99.973625 | 0.001979095 | BGT |
| **McVKB2** | SRR5788131 | 99.98399 | 0.009567606 | BGT |
| **McVKB2** | SRR5788207 | 99.970116 | 0.003958167 | BGT |
| **McVKB2** | SRR5788208 | 99.96208 | 0.005645244 | BGT |
| **McVKB2** | SRR5788209 | 99.96813 | 0.004308748 | BGT |
| **McVKB2** | SRR5788210 | 99.97286 | 0.003741071 | BGT |
| **McVKB2** | SRR5788211 | 99.96458 | 0.005387172 | BGT |
| **McVKB2** | SRR5788212 | 99.97119 | 0.00460325 | BGT |
| **McVKB3** | SRR5788033 | 99.9834 | 0.004746986 | BGT |
| **McVKB3** | SRR5788075 | 99.98971 | 0.001268915 | BGT |
| **McVKB3** | SRR5788026 | 99.98261 | 0.004549774 | BGT |
| **McVKB3** | SRR5788027 | 99.98899 | 0.003357325 | BGT |
| **McVKB3** | SRR5788028 | 99.994484 | 0.001771334 | BGT |
| **McVKB3** | SRR5788030 | 99.99456 | 0.00183901 | BGT |
| **McVKB3** | SRR5788031 | 99.99267 | 0.002535502 | BGT |
| **McVKB3** | SRR5788032 | 99.984024 | 0.004486068 | BGT |
| **McVKB3** | SRR5788130 | 99.973625 | 0.007826659 | BGT |
| **McVKB3** | SRR5788131 | 99.98399 | 0.002860901 | BGT |
| **McVKB3** | SRR5788136 | 99.98759 | 0.004423111 | BGT |
| **McVKB3** | SRR5788137 | 99.98548 | 0.004626535 | BGT |
| **McVKB3** | SRR5788207 | 99.970116 | 0.007138869 | BGT |
| **McVKB3** | SRR5788208 | 99.96208 | 0.009441472 | BGT |
| **McVKB3** | SRR5788209 | 99.96813 | 0.007665759 | BGT |
| **McVKB3** | SRR5788210 | 99.97286 | 0.006719792 | BGT |
| **McVKB3** | SRR5788211 | 99.96458 | 0.009132042 | BGT |
| **McVKB3** | SRR5788212 | 99.97119 | 0.006930989 | BGT |
| **McVSA1** | SRR5720231 | 99.99596 | 0.00404624 | BGT |
| **McVSA1** | SRR5720232 | 99.999 | 0.001003376 | BGT |
| **McVSA1** | SRR5720236 | 99.99674 | 0.003257061 | BGT |
| **McVSA1** | SRR5720237 | 99.99721 | 0.00279628 | BGT |
| **McVSA1** | SRR5720238 | 99.99823 | 0.001772181 | BGT |
| **McVSA1** | SRR5720249 | 99.99895 | 0.001047431 | BGT |
| **McVSA1** | SRR5720250 | 99.99886 | 0.001140056 | BGT |
| **McVSA1** | SRR5720254 | 99.99887 | 0.001135184 | BGT |
| **McVSA1** | SRR5720256 | 99.999146 | 0.000859468 | BGT |
| **McVSA1** | SRR5720259 | 99.99889 | 0.00110498 | BGT |
| **McVSA1** | SRR5788033 | 99.9834 | 0.008805036 | BGT |
| **McVSA1** | SRR5788075 | 99.98971 | 0.001268915 | BGT |
| **McVSA1** | SRR5788089 | 99.99724 | 0.00276746 | BGT |
| **McVSA1** | SRR5788109 | 99.99846 | 0.001543668 | BGT |
| **McVSA1** | SRR5788026 | 99.98261 | 0.012836802 | BGT |
| **McVSA1** | SRR5788027 | 99.98899 | 0.007651798 | BGT |
| **McVSA1** | SRR5788028 | 99.994484 | 0.003742594 | BGT |
| **McVSA1** | SRR5788030 | 99.99456 | 0.003600232 | BGT |
| **McVSA1** | SRR5788031 | 99.99267 | 0.00479299 | BGT |
| **McVSA1** | SRR5788032 | 99.984024 | 0.008878162 | BGT |
| **McVSA1** | SRR5788130 | 99.973625 | 0.016567446 | BGT |
| **McVSA1** | SRR5788131 | 99.98399 | 0.003573253 | BGT |
| **McVSA1** | SRR5788136 | 99.98759 | 0.007988639 | BGT |
| **McVSA1** | SRR5788137 | 99.98548 | 0.00989475 | BGT |
| **McVSA1** | SRR5788207 | 99.970116 | 0.018788047 | BGT |
| **McVSA1** | SRR5788208 | 99.96208 | 0.022836227 | BGT |
| **McVSA1** | SRR5788209 | 99.96813 | 0.019892564 | BGT |
| **McVSA1** | SRR5788210 | 99.97286 | 0.016680054 | BGT |
| **McVSA1** | SRR5788211 | 99.96458 | 0.020902183 | BGT |
| **McVSA1** | SRR5788212 | 99.97119 | 0.01727143 | BGT |
| **McVSA1** | SRR5788229 | 99.99854 | 0.001460913 | BGT |
| **McVSA1** | SRR5788230 | 99.998886 | 0.001112999 | BGT |
| **McVSA1** | SRR5788233 | 99.99854 | 0.001451874 | BGT |
| **McVSA1** | SRR5788283 | 99.99851 | 0.001486994 | BGT |
| **McVSA1** | SRR5788282 | 99.99712 | 0.002877038 | BGT |
| **McVSA1** | SRR5788281 | 99.996635 | 0.003369628 | BGT |
| **McVSA1** | SRR5788284 | 99.99817 | 0.001829551 | BGT |
| **McVSA1** | SRR5788285 | 99.99818 | 0.001823641 | BGT |
| **McVSA1** | SRR5788286 | 99.99813 | 0.001866522 | BGT |
| **McVSA1** | SRR5788288 | 99.998695 | 0.001302452 | BGT |
| **McVSA1** | SRR5788318 | 99.9988 | 0.001199123 | BGT |
| **McVSA1** | SRR5788374 | 99.99895 | 0.001055628 | BGT |
| **McVSA1** | SRR5788430 | 99.99832 | 0.001682257 | BGT |
| **McVSA1** | SRR5788431 | 99.99872 | 0.001279427 | BGT |
| **McVSA1** | SRR5788436 | 99.99881 | 0.00118995 | BGT |
| **McVSA1** | SRR5788435 | 99.998566 | 0.001428942 | BGT |
| **McVSA1** | SRR5788429 | 99.998055 | 0.00194779 | BGT |
| **McVSA1** | SRR9178106 | 99.99787 | 0.002129238 | Mende |
| **McVSA1** | SRR9178213 | 99.998184 | 0.001818341 | Mende |
| **McVSA1** | SRR9178292 | 99.99803 | 0.001966518 | Mende |
| **McVSA1** | SRR9178368 | 99.99813 | 0.001863446 | Mende |
| **McVSA1** | SRR9178358 | 99.998146 | 0.001852308 | Mende |
| **McVSA1** | SRR9178335 | 99.997986 | 0.002012756 | Mende |
| **McVSA1** | SRR9178205 | 99.99348 | 0.006520916 | Mende |
| **McVSA1** | SRR9178320 | 99.994 | 0.005995357 | Mende |
| **McVKB2** | ERR594376 | 99.99707 | 0.002930175 | Tara |
| **McVKB2** | ERR594392 | 99.965965 | 0.000689605 | Tara |
| **McVKB2** | ERR594414 | 99.921646 | 0.002198584 | Tara |
| **McVKB2** | ERR594382 | 99.92059 | 0.002130446 | Tara |
| **McVKB2** | ERR594361 | 99.9567 | 0.0013392 | Tara |
| **McVKB2** | ERR594359 | 99.95748 | 0.001332111 | Tara |
| **McVKB2** | ERR594389 | 99.98083 | 0.001331401 | Tara |
| **McVKB2** | ERR594362 | 99.98643 | 0.00087922 | Tara |
| **McVKB2** | ERR599350 | 99.99125 | 0.003240518 | Tara |
| **McVKB3** | ERR594385 | 99.995705 | 0.001500044 | Tara |
| **McVKB3** | ERR594392 | 99.965965 | 0.020873025 | Tara |
| **McVKB3** | ERR594414 | 99.921646 | 0.04176986 | Tara |
| **McVKB3** | ERR594382 | 99.92059 | 0.042290036 | Tara |
| **McVKB3** | ERR594361 | 99.9567 | 0.025936551 | Tara |
| **McVKB3** | ERR594359 | 99.95748 | 0.02555807 | Tara |
| **McVKB3** | ERR594389 | 99.98083 | 0.013581123 | Tara |
| **McVKB3** | ERR594362 | 99.98643 | 0.009690655 | Tara |
| **McVKB3** | ERR599374 | 99.99579 | 0.00303531 | Tara |
| **McVKB4** | ERR594414 | 99.921646 | 0.003272586 | Tara |
| **McVKB4** | ERR594382 | 99.92059 | 0.003310048 | Tara |
| **McVKB4** | ERR599350 | 99.99125 | 0.002187865 | Tara |
| **McVSA1** | ERR594385 | 99.995705 | 0.002795147 | Tara |
| **McVSA1** | ERR594392 | 99.965965 | 0.012472245 | Tara |
| **McVSA1** | ERR594414 | 99.921646 | 0.031116072 | Tara |
| **McVSA1** | ERR594382 | 99.92059 | 0.03167623 | Tara |
| **McVSA1** | ERR594361 | 99.9567 | 0.016018566 | Tara |
| **McVSA1** | ERR594359 | 99.95748 | 0.01562878 | Tara |
| **McVSA1** | ERR594389 | 99.98083 | 0.004261006 | Tara |
| **McVSA1** | ERR594362 | 99.98643 | 0.003005065 | Tara |
| **McVSA1** | ERR599350 | 99.99125 | 0.003320088 | Tara |
| **McVSA1** | ERR599374 | 99.99579 | 0.001176947 | Tara |

| Genome | SRA/ENA_Accession | unmapped(%) | RelativeAbundance(%) | DataSet |
| --- | --- | --- | --- | --- |
| McVKB2 | SRR5788033 | 99.9834 | 0.00304499 | BGT |
| McVKB2 | SRR5788075 | 99.98971 | 0.00126892 | BGT |
| McVKB2 | SRR5788032 | 99.984024 | 0.0026116 | BGT |
| McVKB2 | SRR5788130 | 99.973625 | 0.0019791 | BGT |
| McVKB2 | SRR5788131 | 99.98399 | 0.00956761 | BGT |
| McVKB2 | SRR5788207 | 99.970116 | 0.00395817 | BGT |
| McVKB2 | SRR5788208 | 99.96208 | 0.00564524 | BGT |
| McVKB2 | SRR5788209 | 99.96813 | 0.00430875 | BGT |
| McVKB2 | SRR5788210 | 99.97286 | 0.00374107 | BGT |
| McVKB2 | SRR5788211 | 99.96458 | 0.00538717 | BGT |
| McVKB2 | SRR5788212 | 99.97119 | 0.00460325 | BGT |
| McVKB2 | ERR594376 | 99.99707 | 0.00293018 | Tara |
| McVKB2 | ERR594392 | 99.965965 | 0.00068961 | Tara |
| McVKB2 | ERR594414 | 99.921646 | 0.00219858 | Tara |
| McVKB2 | ERR594382 | 99.92059 | 0.00213045 | Tara |
| McVKB2 | ERR594361 | 99.9567 | 0.0013392 | Tara |
| McVKB2 | ERR594359 | 99.95748 | 0.00133211 | Tara |
| McVKB2 | ERR594389 | 99.98083 | 0.0013314 | Tara |
| McVKB2 | ERR594362 | 99.98643 | 0.00087922 | Tara |
| McVKB2 | ERR599350 | 99.99125 | 0.00324052 | Tara |
| McVKB3 | SRR5788033 | 99.9834 | 0.00474699 | BGT |
| McVKB3 | SRR5788075 | 99.98971 | 0.00126892 | BGT |
| McVKB3 | SRR5788026 | 99.98261 | 0.00454977 | BGT |
| McVKB3 | SRR5788027 | 99.98899 | 0.00335733 | BGT |
| McVKB3 | SRR5788028 | 99.994484 | 0.00177133 | BGT |
| McVKB3 | SRR5788030 | 99.99456 | 0.00183901 | BGT |
| McVKB3 | SRR5788031 | 99.99267 | 0.0025355 | BGT |
| McVKB3 | SRR5788032 | 99.984024 | 0.00448607 | BGT |
| McVKB3 | SRR5788130 | 99.973625 | 0.00782666 | BGT |
| McVKB3 | SRR5788131 | 99.98399 | 0.0028609 | BGT |
| McVKB3 | SRR5788136 | 99.98759 | 0.00442311 | BGT |
| McVKB3 | SRR5788137 | 99.98548 | 0.00462654 | BGT |
| McVKB3 | SRR5788207 | 99.970116 | 0.00713887 | BGT |
| McVKB3 | SRR5788208 | 99.96208 | 0.00944147 | BGT |
| McVKB3 | SRR5788209 | 99.96813 | 0.00766576 | BGT |
| McVKB3 | SRR5788210 | 99.97286 | 0.00671979 | BGT |
| McVKB3 | SRR5788211 | 99.96458 | 0.00913204 | BGT |
| **McVKB3** | SRR5788212 | 99.97119 | 0.00693099 | BGT |
| **McVKB3** | ERR594385 | 99.995705 | 0.00150004 | Tara |
| **McVKB3** | ERR594392 | 99.965965 | 0.02087303 | Tara |
| **McVKB3** | ERR594414 | 99.921646 | 0.04176986 | Tara |
| **McVKB3** | ERR594382 | 99.92059 | 0.04229004 | Tara |
| **McVKB3** | ERR594361 | 99.9567 | 0.02593655 | Tara |
| **McVKB3** | ERR594359 | 99.95748 | 0.02555807 | Tara |
| **McVKB3** | ERR594389 | 99.98083 | 0.01358112 | Tara |
| **McVKB3** | ERR594362 | 99.98643 | 0.00969066 | Tara |
| **McVKB3** | ERR599374 | 99.99579 | 0.00303531 | Tara |
| **McVKB4** | ERR594414 | 99.921646 | 0.00327259 | Tara |
| **McVKB4** | ERR594382 | 99.92059 | 0.00331005 | Tara |
| **McVKB4** | ERR599350 | 99.99125 | 0.00218787 | Tara |
| **McVSA1** | SRR5720231 | 99.99596 | 0.00404624 | BGT |
| **McVSA1** | SRR5720232 | 99.999 | 0.00100338 | BGT |
| **McVSA1** | SRR5720236 | 99.99674 | 0.00325706 | BGT |
| **McVSA1** | SRR5720237 | 99.99721 | 0.00279628 | BGT |
| **McVSA1** | SRR5720238 | 99.99823 | 0.00177218 | BGT |
| **McVSA1** | SRR5720249 | 99.99895 | 0.00104743 | BGT |
| **McVSA1** | SRR5720250 | 99.99886 | 0.00114006 | BGT |
| **McVSA1** | SRR5720254 | 99.99887 | 0.00113518 | BGT |
| **McVSA1** | SRR5720256 | 99.999146 | 0.00085947 | BGT |
| **McVSA1** | SRR5720259 | 99.99889 | 0.00110498 | BGT |
| **McVSA1** | SRR5788033 | 99.9834 | 0.00880504 | BGT |
| **McVSA1** | SRR5788075 | 99.98971 | 0.00126892 | BGT |
| **McVSA1** | SRR5788089 | 99.99724 | 0.00276746 | BGT |
| **McVSA1** | SRR5788109 | 99.99846 | 0.00154367 | BGT |
| **McVSA1** | SRR5788026 | 99.98261 | 0.0128368 | BGT |
| **McVSA1** | SRR5788027 | 99.98899 | 0.0076518 | BGT |
| **McVSA1** | SRR5788028 | 99.994484 | 0.00374259 | BGT |
| **McVSA1** | SRR5788030 | 99.99456 | 0.00360023 | BGT |
| **McVSA1** | SRR5788031 | 99.99267 | 0.00479299 | BGT |
| **McVSA1** | SRR5788032 | 99.984024 | 0.00887816 | BGT |
| **McVSA1** | SRR5788130 | 99.973625 | 0.01656745 | BGT |
| **McVSA1** | SRR5788131 | 99.98399 | 0.00357325 | BGT |
| **McVSA1** | SRR5788136 | 99.98759 | 0.00798864 | BGT |
| **McVSA1** | SRR5788137 | 99.98548 | 0.00989475 | BGT |
| **McVSA1** | SRR5788207 | 99.970116 | 0.01878805 | BGT |
| **McVSA1** | SRR5788208 | 99.96208 | 0.02283623 | BGT |
| McVSA1 | SRR5788209 | 99.96813 | 0.01989256 | BGT |
| McVSA1 | SRR5788210 | 99.97286 | 0.01668005 | BGT |
| McVSA1 | SRR5788211 | 99.96458 | 0.02090218 | BGT |
| McVSA1 | SRR5788212 | 99.97119 | 0.01727143 | BGT |
| McVSA1 | SRR5788229 | 99.99854 | 0.00146091 | BGT |
| McVSA1 | SRR5788230 | 99.998886 | 0.001113 | BGT |
| McVSA1 | SRR5788233 | 99.99854 | 0.00145187 | BGT |
| McVSA1 | SRR5788283 | 99.99851 | 0.00148699 | BGT |
| McVSA1 | SRR5788282 | 99.99712 | 0.00287704 | BGT |
| McVSA1 | SRR5788281 | 99.996635 | 0.00336963 | BGT |
| McVSA1 | SRR5788284 | 99.99817 | 0.00182955 | BGT |
| McVSA1 | SRR5788285 | 99.99818 | 0.00182364 | BGT |
| McVSA1 | SRR5788286 | 99.99813 | 0.00186652 | BGT |
| McVSA1 | SRR5788288 | 99.998695 | 0.00130245 | BGT |
| McVSA1 | SRR5788318 | 99.9988 | 0.00119912 | BGT |
| McVSA1 | SRR5788374 | 99.99895 | 0.00105563 | BGT |
| McVSA1 | SRR5788430 | 99.99832 | 0.00168226 | BGT |
| McVSA1 | SRR5788431 | 99.99872 | 0.00127943 | BGT |
| McVSA1 | SRR5788436 | 99.99881 | 0.00118995 | BGT |
| McVSA1 | SRR5788435 | 99.998566 | 0.00142894 | BGT |
| McVSA1 | SRR5788429 | 99.998055 | 0.00194779 | BGT |
| McVSA1 | SRR9178106 | 99.99787 | 0.00212924 | HOT |
| McVSA1 | SRR9178213 | 99.998184 | 0.00181834 | HOT |
| McVSA1 | SRR9178292 | 99.99803 | 0.00196652 | HOT |
| McVSA1 | SRR9178368 | 99.99813 | 0.00186345 | HOT |
| McVSA1 | SRR9178358 | 99.998146 | 0.00185231 | HOT |
| McVSA1 | SRR9178335 | 99.997986 | 0.00201276 | HOT |
| McVSA1 | SRR9178205 | 99.99348 | 0.00652092 | HOT |
| McVSA1 | SRR9178320 | 99.994 | 0.00599536 | HOT |
| McVSA1 | ERR594385 | 99.995705 | 0.00279515 | Tara |
| McVSA1 | ERR594392 | 99.965965 | 0.01247225 | Tara |
| McVSA1 | ERR594414 | 99.921646 | 0.03111607 | Tara |
| McVSA1 | ERR594382 | 99.92059 | 0.03167623 | Tara |
| McVSA1 | ERR594361 | 99.9567 | 0.01601857 | Tara |
| McVSA1 | ERR594359 | 99.95748 | 0.01562878 | Tara |
| McVSA1 | ERR594389 | 99.98083 | 0.00426101 | Tara |
| McVSA1 | ERR594362 | 99.98643 | 0.00300507 | Tara |
| McVSA1 | ERR599350 | 99.99125 | 0.00332009 | Tara |
| McVSA1 | ERR599374 | 99.99579 | 0.00117695 | Tara |

**Supplementary Figure S1.**
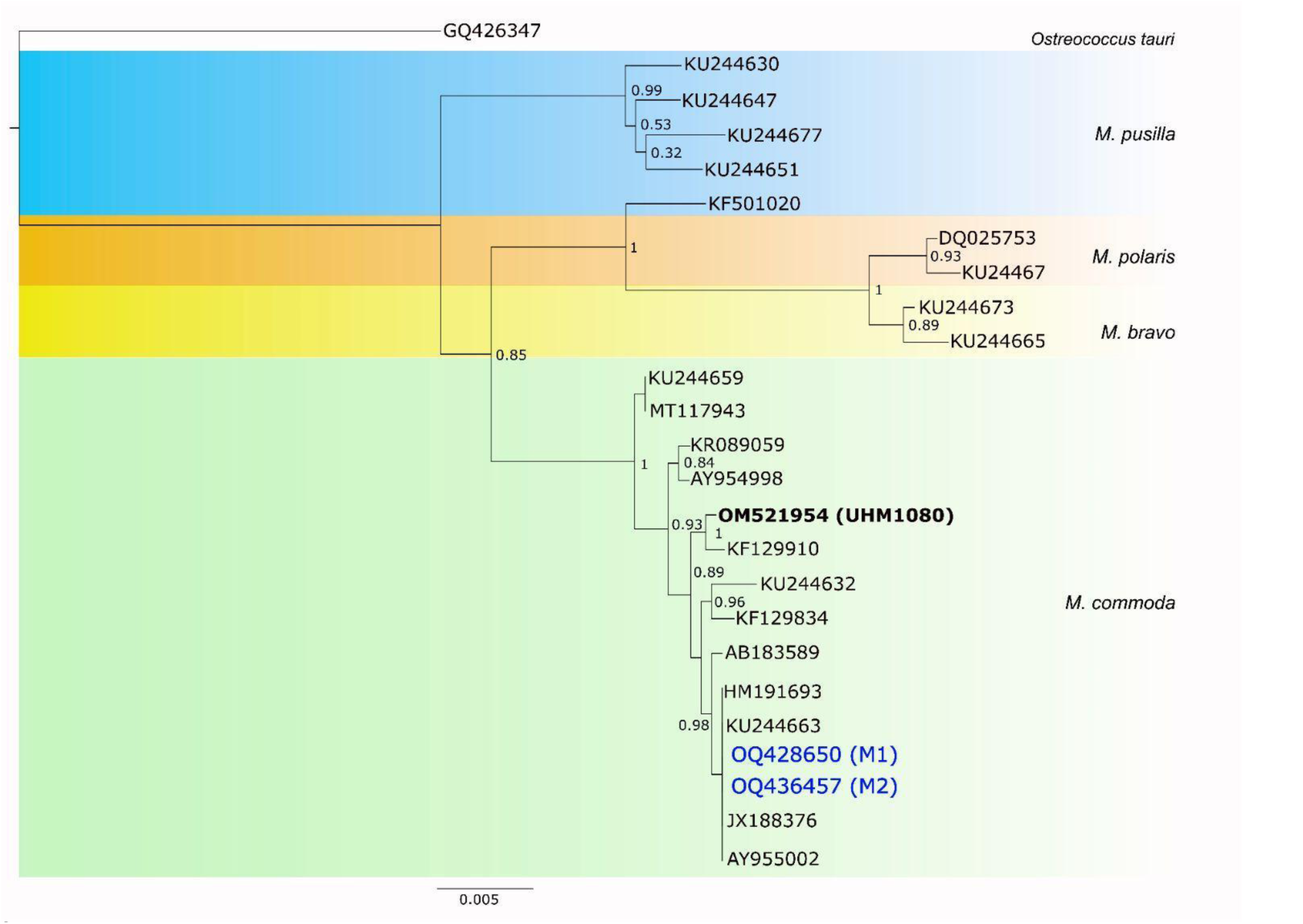
A phylogenetic tree of Mamiellales. The tree is based on trimmed alignments of partial 18S rRNA genes from the two *Micromonas* strains used in this study (M1 and M2), a *Micromonas* strain from the pelagic Station ALOHA (UHM1080), and related sequences found using NCBI’s BLAST tool. Alignments were created and trimmed with Geneious 11.1 default alignment tool and processed through FastTree using approximately-maximum-likelihood. Node support values reflect FastTree local support values derived from the Shimodaira-Hasegawa test. Figure reproduced with permission from Bedi de Silva et al. (2024) Environ. Microbiol 26(8) e16686,

**Supplementary Figure S2.**
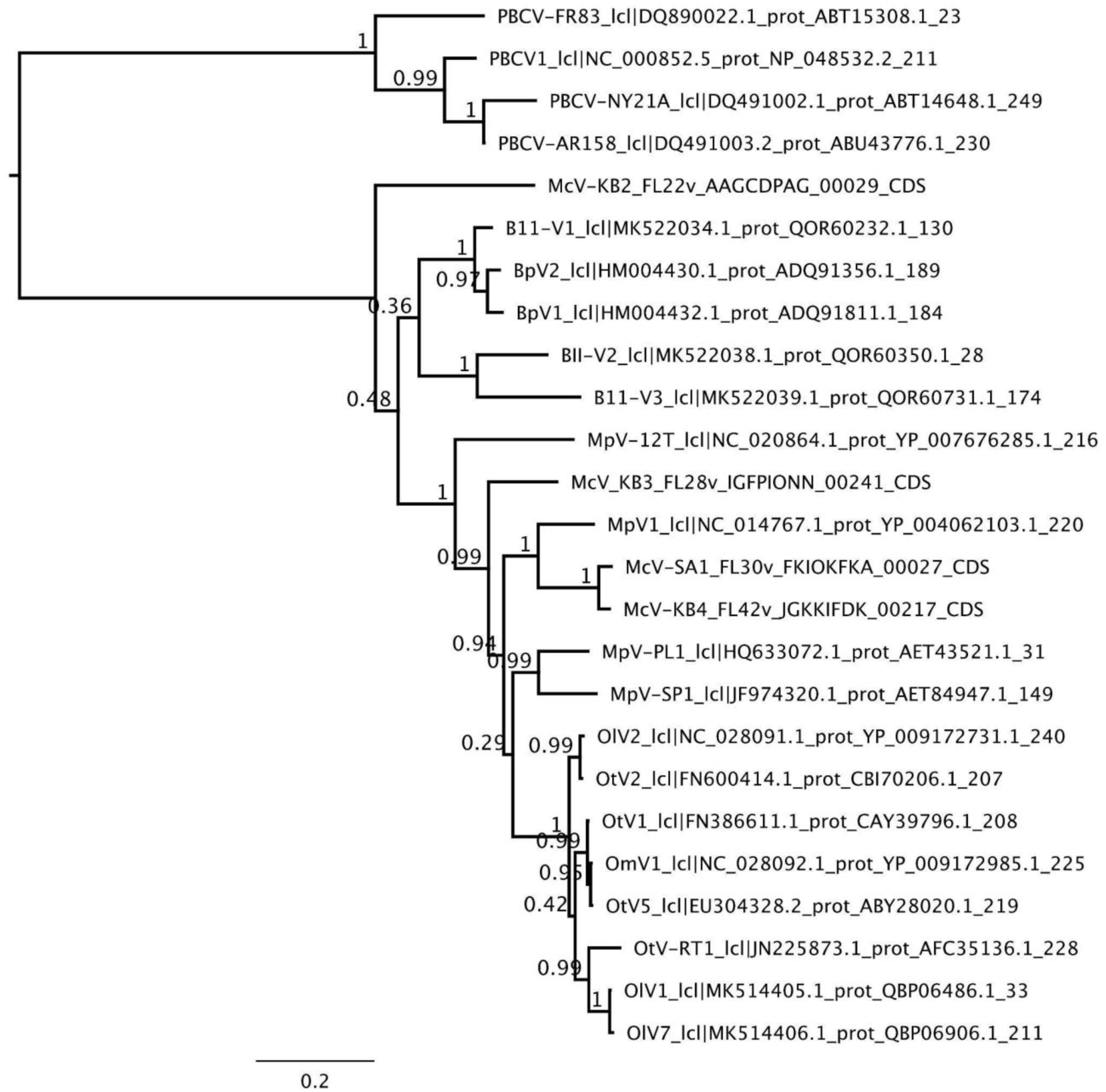
Prasinovirus and chlorovirus species tree based on the polB orthogroup. Tree was created using FastTree, scale bar represents substitutions per site.

**Supplementary Figure S3.**
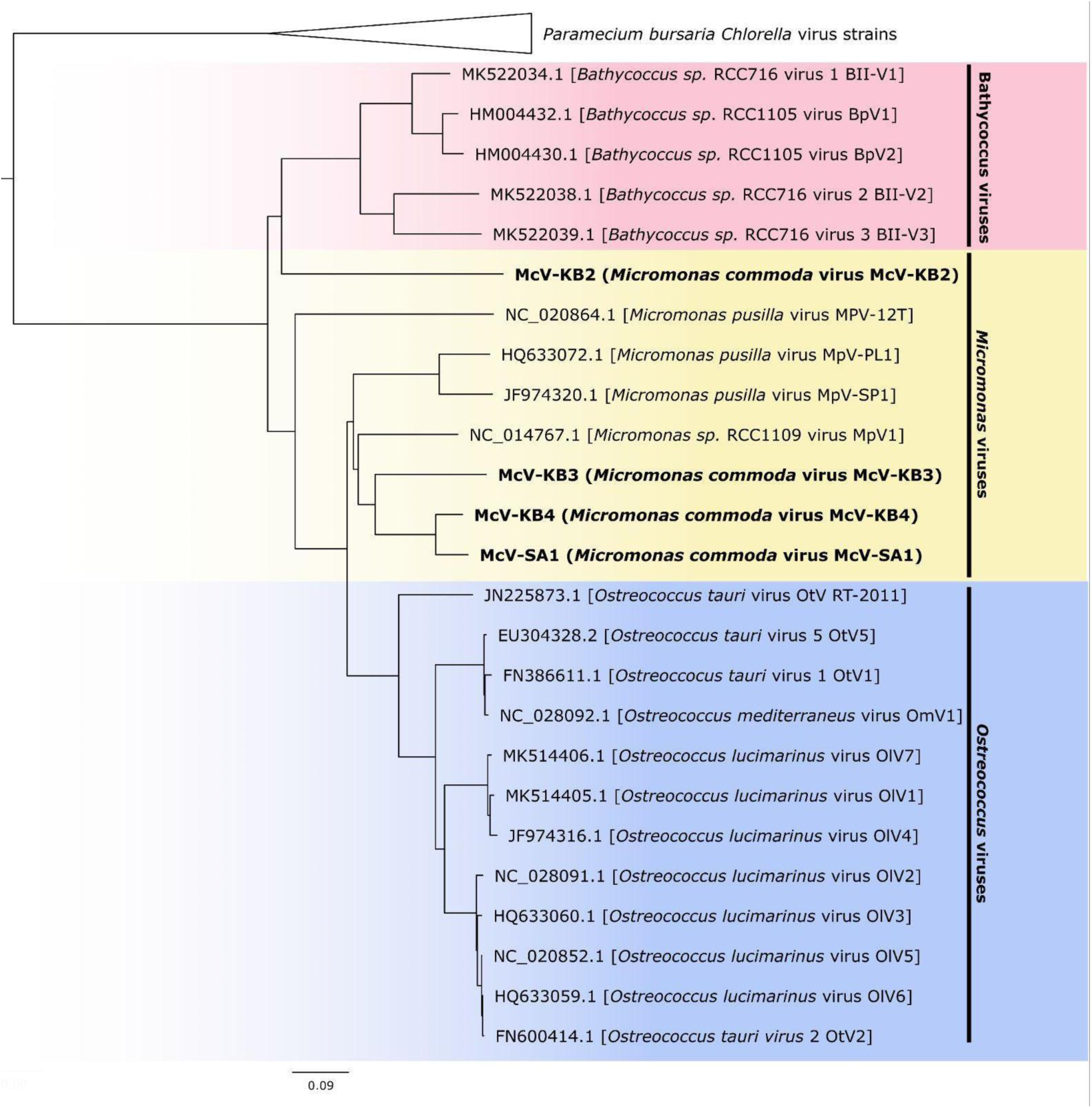
STAG-generated species tree using the 26 orthogroups that are possessed by all prasinoviruses and chloroviruses genomes used in our OrthoFinder analysis. STAG bipartition support values are not available for datasets with fewer than 100 shared orthogroups. Scale bar indicates substitutions per site.

**Supplementary Figure S4.**
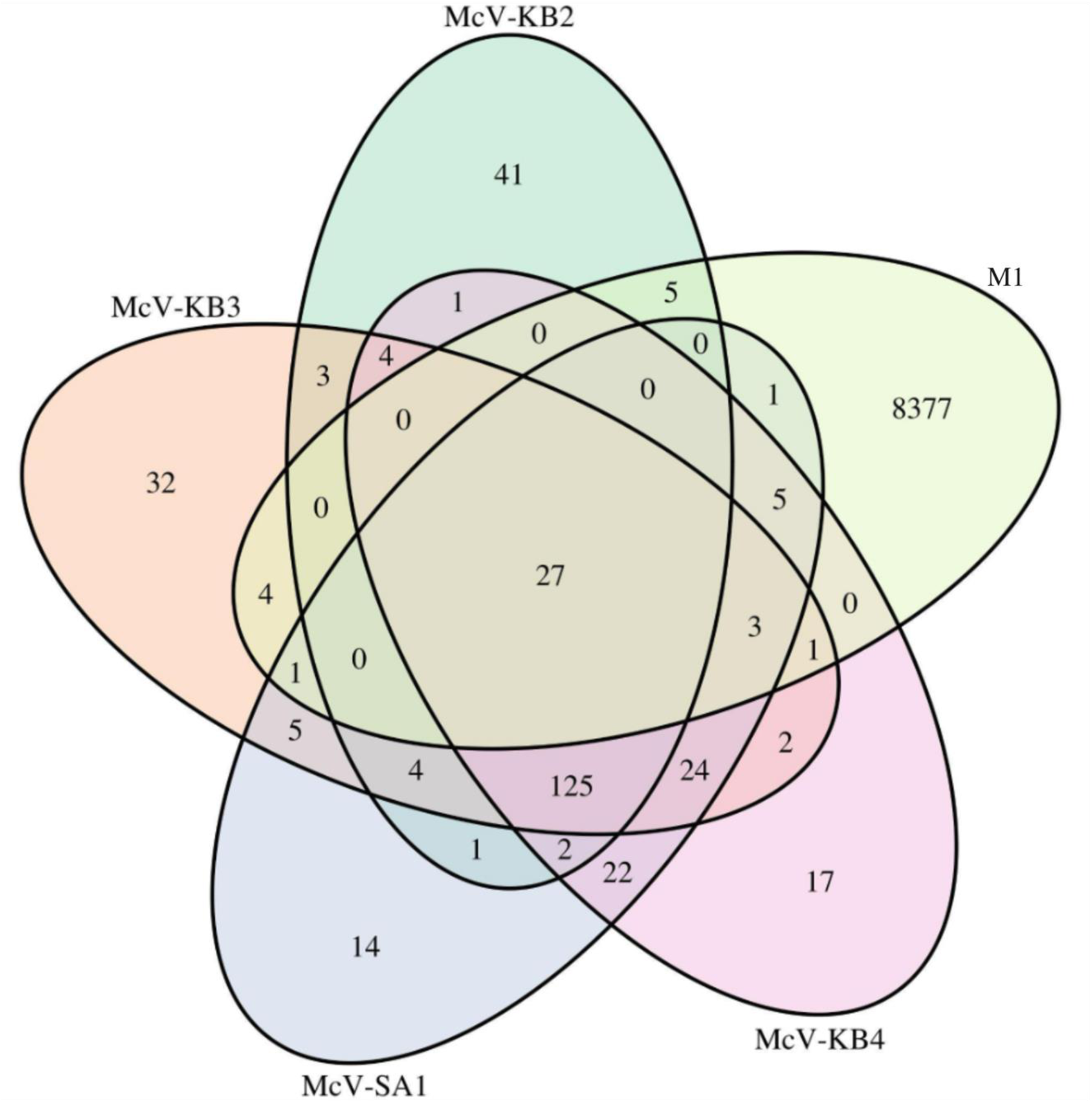
Venn Diagram of the number of orthogroups shared between the four HiMcVs and host M1.

**Supplementary Figure S5.**
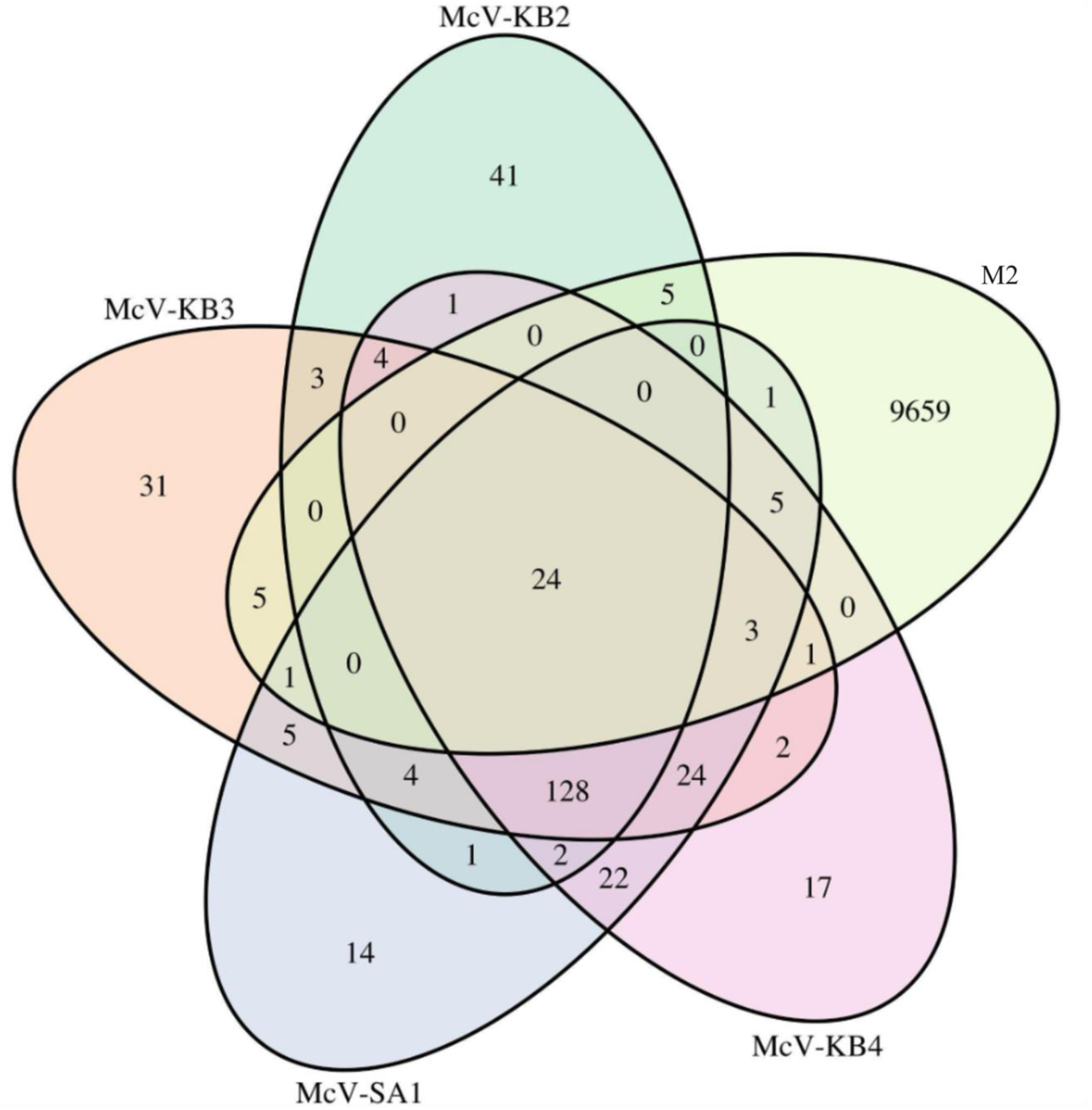
Venn Diagram of the number of orthogroups shared between the four HiMcVs and host M2. Clustering analysis was conducted with OrthoFinder.

**Supplementary Figure S6.**
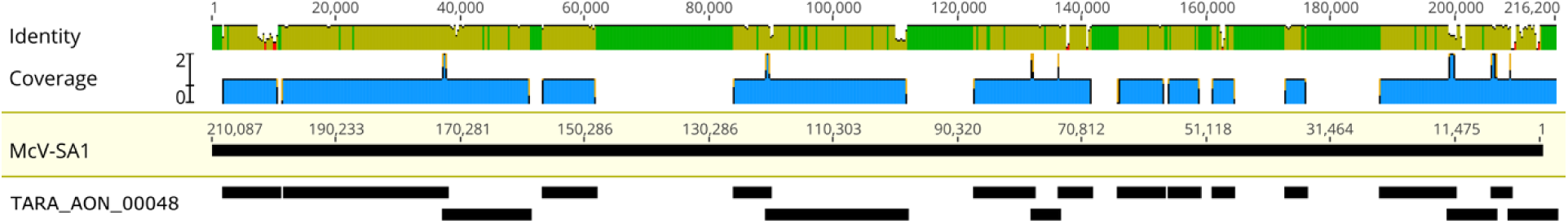
Contigs from GVMAG TARA_AON_NCLDV_00048 mapped to McV-SA1. Average coverage was 71%, with an average 92% nucleotide identity. Mapping of contigs was conducted in Geneious 11.1.

